# A single E205D allele of a key P450 *CYP6P3* is driving metabolic pyrethroid resistance in the major African malaria vector *Anopheles gambiae*

**DOI:** 10.1101/2024.02.18.580859

**Authors:** Jonas A. Kengne-Ouafo, Mersimine Kouamo, Abdullahi Muhammad, Arnaud Tepa, Stevia Ntadoun, Leon Mugenzi, Theofelix Tekoh, Jack Hearn, Magellan Tchouakui, Murielle Wondji, Sulaiman S. Ibrahim, Charles S. Wondji

## Abstract

Deciphering the molecular drivers of insecticide resistance is paramount to extend the effectiveness of malaria vector control tools. Here, we demonstrated that the E205D amino acid change in a key metabolic resistance P450 *CYP6P3* drives pyrethroid resistance in the major malaria vector, *Anopheles gambiae*. Spatio-temporal whole genome Poolseq analyses in Cameroon detected a major P450-linked locus on chromosome 2R beside the sodium channel locus. *In vitro* metabolism assays with recombinantly expressed *CYP6P3* protein revealed that the catalytic efficiency of 205D was 2.5 times higher than E205 with α-cypermethrin. Similar patterns were observed for permethrin. Overexpression of the 205D allele in transgenic flies confers higher more pyrethroids and carbamates resistance, compared to controls. A DNA-based assay further supported that the *CYP6P3*-205D variant strongly correlates with pyrethroid resistance in field populations (OR=26.4; P<0.0001) and that it reduces the efficacy of pyrethroid-only LLINs with homozygote RR genotype exhibiting significantly higher survival following PermaNet 3.0 exposure compared to the SS genotype (OR: 6.1, p = 0.0113). Furthermore, the *CYP6P3*-E205D combines with the *kdr* target-site resistance mechanisms to worsen the loss of bednet efficacy. The 205D mutation is now predominant in West and Central Africa but less abundant or absent in East and South Africa with signs of introgression with *An. coluzzii* in Ghana. This study highlights the importance of P450-based resistance and designs field-applicable tools to easily track the spread of metabolic resistance and assess its impact on control interventions.

**One Sentence Summary:** The major obstacle to malaria control and elimination is the spread of parasite resistance to anti-malarial drugs, and mosquito resistance to insecticides. In this study, we identified a key point mutation E205D in the metabolic gene *CYP6P3* (cytochrome P450) conferring resistance to pyrethroids by enhancing the breakdown of insecticides used for bednets impregnation. DNA-based assays were then designed and used to determine the spread of the resistance across Africa and demonstrate that the *CYP6P3*-205D allele works together with the knockdown resistance in the voltage-gated Sodium channel to reduce the efficacy of insecticide-treated bednets.

## Introduction

Malaria burden remains high with more than 600,000 and 200 million yearly recorded deaths and cases respectively worldwide (*1*). The impact of the disease is more felt in Sub-Saharan Africa which alone experiences more than 90% of the burden. Progress made since 2000 has stalled resulting in the flattening of the previously decreasing course of malaria morbidity and mortality (*2*) with resistance, especially towards the pyrethroids as one of the major factors for the lack of progress (*3–5*). This has been shown by several randomised control trials revealing a greater reduction of malaria prevalence when using the next-generation long-lasting insecticidal bed nets (LLINs), such as the PBO-nets (*6*) or recent Interceptor G2 nets (*7*).

Insecticide resistance should be tackled to improve the effectiveness of current and future tools (*8*). However, understanding the underlying resistance mechanisms is a key prerequisite to help design suitable resistance management strategies. The main resistance mechanisms are target site and metabolic resistance (*8*). If progress has been made for *kdr* (*9*) with makers detected back in 1998, metabolic resistance has been more complex. Although genes conferring pyrethroid resistance have been detected and functionally characterised in many malaria vector species including the *An. gambiae* (*CYP6P3*, *CYP6M2,* etc) and *An. funestus* (*CYP6P9a*, *CYP6P9b*, etc), it was only recently that a DNA-based marker was detected for a P450-based resistance in *An. funestus* (*10, 11*). For *An. gambiae*, studies aiming at detecting genetic variants extensively associated with pyrethroid resistance (*12–16*) through modulation of metabolic activity (*17, 18*) are limited. This lack of DNA-based assay has limited the detection of metabolic resistance in the major vector *An. gambiae* and prevented establishing the extent to which this resistance mechanism is reducing efficacy of existing tools. The recent report of one such marker in the *CYP6P4* gene in East Africa is a strong indication that the success made in *An. funestus* could be achieved also in *An. gambiae* (*19*).

Previous efforts to detecting genetic variants associated with pyrethroid resistance (*16*) have relied on whole genome sequencing of wild mosquitoes which has helped detect strong signature of selection Africa-wide (*16*). However, it has been more challenging to clearly detect variants associated with resistance. One reason of this challenge is the high level of resistance already in the field making it difficult to associate a variant to the resistance phenotypes. This has led to the strategy using hybrid strain generated from highly resistant mosquitoes and laboratory susceptible colonies to segregate the variant and the resistance phenotypes as done in *Ae. aegypti* previously (*20*) and *An funestus* (*11*)

Here using a whole genome sequencing approach and hybrid strains, we detected a strong signal of selection around the P450 cluster on 2R containing the *CYP6P3* gene. A single amino acid change E205D was shown to be tightly linked to resistance allowing the design of a DNA-based tool for *CYP6P3*-linked pyrethroid resistance. Using this assay, we revealed the contribution of E205D variant to type I and Type II pyrethroid resistance, its geographical distribution and temporal evolution, as well as its role in reduced efficacy to LLINs. Moreover, the combined effect of this P450-based resistance and *kdr* mutation was established.

## Results

### Creation of hybrid mosquito lines, resistance phenotyping and *kdr* genotyping

Crossing was carried out to ensure the segregation of resistance genotypes in the strain. A total of 150 female of susceptible mosquitoes (Kisumu colonies) were individually crossed with 300 male resistant mosquitoes from the field (Mangoum, 2022). WHO tube bioassay with 0.75% (1X) permethrin revealed mortality rates of 2.5% and 100% for Mangoum_2022 and Kisumu mosquitoes, respectively prior to crossing. After crossing, the mortality assessment at generations F_1_, F_3_ and F_4_ showed a stable level of resistance with mortality to 1X permethrin around 76% after 1 h exposure (Fig. S1 **A**). The *kdr*-1014F mutant allele was fixed in the field population (Mangoum_2022) and absent in Kisumu. However, all the genotypes were found in the F_4_ hybrid mosquitoes in the following proportions 17.5%, 32.5% and 50% for the RR, RS and SS genotypes respectively attesting genotypes re-segregation. (Fig. S1 **B**). For the selection of highly susceptible and highly resistant mosquitoes for the molecular work, F_4_ hybrids were subjected to different insecticides at the corresponding LT_20s_ and LT_80s_ as indicated below.

### Whole genome Pooled DNA sequencing (PoolSeq)

Sequencing of phenotyped hybrid mosquitoes (pooled DNA samples) was performed to detect genetic variants associated with pyrethroid resistance. To achieve that, a total of 10 DNA pools made up of 50 mosquitoes each were sequenced among which 3 pools were obtained from hybrid mosquitoes that survived 90 min (LT_80_) permethrin exposure (resistant/alive), 3 pools from hybrid mosquitoes that died after 3 min (LT_20_) permethrin exposure (susceptible/dead), 2 pools of unexposed Kisumu lab colony, and 2 pools of unexposed F_0_ Mangoum mosquitoes collected separately in 2018 and 2022. Alignment of trimmed raw reads from pool sequencing against the *Anopheles gambiae* reference genome version 59 (VectorBase.org) was done successfully with the percentage mapped reads ranging from 74% for field mosquitoes to 98.6% for hybrid and lab mosquitoes (Table S2) and mean coverage between 31% and 66%. Principal component analysis (PCA) revealed clustering of samples according to origin and phenotype. The 3 replicates of alive hybrid mosquitoes grouped together away from the dead and Kisumu groups (Fig. S2 **A**). The same pattern was observed after merging the replicates of each group (Fig. S2 **B**), with the Alive group clustering separately from the Dead and Kisumu, which cluster together, and closer to the Mangoum field samples.

### PoolSeq WGAS detects major genomic loci associated with permethrin resistance

To identify genomic regions associated with resistance, we compared resistant field mosquitoes (Mangoum 2018 and Mangoum 2022) and the F_4_ Kisumu_X_Mangoum permethrin-alive mosquitoes (resistant hybrids) to the susceptible Kisumu colony and F_4_ Kisumu_X_Mangoum permethrin-dead mosquitoes (susceptible hybrids) using the fixation index estimate (F_st_). Similar peaks were obtained upon comparing resistant field mosquitoes to Kisumu and susceptible hybrids (Fig. 1 **A** & **B**; Fig. S3 **A** & **B**). The main peaks were located on chromosome 2L in the regions covering the voltage-gated sodium channel (VGSC) (F_st_ = 0.401), short chain dehydrogenase (AGAP005167) (F_st_ = 0.23) and LIM homeobox protein (AGAP006540) (F_st_ = 0.211). On chromosome 2R the loci of interest were those spanning the detoxifying CYP6 cluster with a F_st_ peak of 0.225, the serine protein kinase region (AGAP001683) (F_st_ = 0.213), ubiquitin carboxyl-terminal hydrolase 36 region (AGAP003643, (F_st_ = 0.231), this region/peak also contained the keap 19 gene (AGAP003645) and the putative gastrin/bombesin receptor 2 gene (AGAP003631). The last peak in this chromosome with a Fst max of 0.258 was found within the region containing the salivary gland protein 5 (AGAP004334). This region also covers the detoxifying GSTD cluster and some odorant receptor genes (AGAP004278, AGAP004278), a synaptic vesicle protein (AGAP004354) and a CLIP-domain serine protease (AGAP004317). The other top peaks on this chromosome were mainly within genes with unknown functions (unspecified products). Another cluster of detoxification genes (glutathione-S-transferase Epsilon (GSTE) was prominent on chromosome 3R with Fst max of 0.403. The two main peaks found on chromosome X were within the region containing the NADPH cytochrome reductase (*CPR*, Fst = 0.220) and the cytochrome P450, *CYP9K1*(Fst = 0.219) respectively (Fig. 1 **A** & **B**; Fig. S3 **A** & **B**). The *GSTT2* (AGAP000888) gene was also found around the region (Fst = 0.168). However, peaks in chromosomes 2L and 2R were abrogated when Fst was computed between resistant field and resistant hybrid mosquitoes suggesting those signals may be playing a role in resistance to permethrin in both mosquito lines (Fig S3. **c**). The VGSC peak considerably reduced in magnitude but not completely (from Fst of 0.401 to 0.142) implying higher frequency of the involved mutations in field mosquitoes. Worth noting was the complete absence/abrogation of signals in chromosomes 3L, 3R and X up on comparing resistant hybrids to their susceptible counterparts (Fig. 1 **B**). Though at lower magnitude, previously seen signals between field vs Kisumu in chromosomes 2L and 2R were still present (they reduced from Fst of 0.401to 0.084 and from Fst of 0.225 to 0.073 for VGSC and CYP6 regions respectively) (Fig. 1 **B**) emphasizing the fact that those might be the key signals involved in permethrin resistance. To confirm these signals, we carried out the Cochran-Mantel-Haenszel (CMH) test which is a GWAS-like test that detects consistency in allele frequency changes in biological replicates. The CMH test validated the signals in chromosome 2L, 2R and UNKN (Fig. 1 **C**). P values as low as 8.11e-13 and 1.04e-12 were obtained in VGSC and CYP6 regions respectively. To further understand the data, nucleotide diversity and signs of selection were assessed within a 300kb region covering the signals. The region was also found to be likely under a positive selection as indicated by the negative Tajima’s D values obtained for field resistant samples ranging from -1.41 to -0.831. Kisumu and susceptible hybrid mosquitoes had values close to zero indicating neutrality whereas the resistant hybrid population had relatively positive values ranging from 0.278 to 0.748 (Fig. 1 **D**). A marked reduction in nucleotide diversity was observed in the CYP6 cluster in resistant field samples (Mangoum 2018 and 2022) with mean pi of 0.0028 and 0.0074, respectively compared to 0.0095 and 0.013 in susceptible dead mosquitoes and the reference Kisumu respectively. The resistant hybrid mosquitoes were also the most diverse in that region with a mean pi of 0.0138 (Fig. 1 **E**). This could be attributed to the lab conditions and short propagation period (4 generations only not enough for ideal recombination to occur). Further, zooming within the CYP6 cluster (10kb) revealed a pronounced reduced diversity in *CYP6P3* genes and the pseudo gene *CYP6P1* (Fig. 1 **F**). For the short chain dehydrogenase cluster in the 2L chromosome, the nucleotide diversity was low in the susceptible populations and high in the resistant ones. The nucleotide diversity was globally low in VGSC cluster for all populations except the resistant hybrid mosquitoes. Apart from the CYP6 cluster, no other cluster showed a sign of selection (Fig S4). Therefore, we focused on this locus to identify the candidate variants associated with metabolic resistance to pyrethroid in these populations of *An. gambiae*.

**Figure 1:**
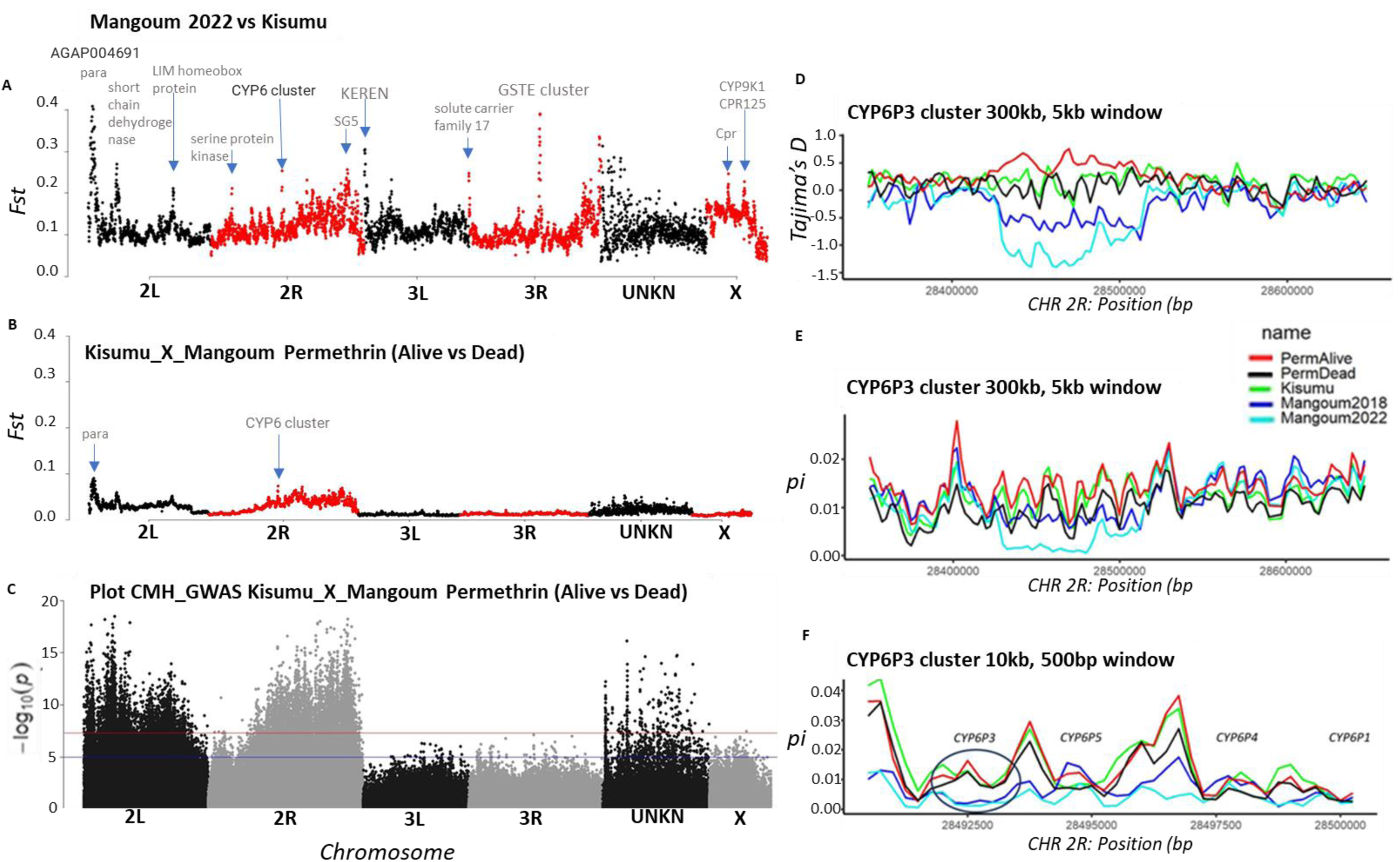
Detection of CYP6P3 loci. (A) Pair-wise Fst comparison between Fst Mangoum 2022 and Kisumu. (B) Fst F4 Kisumu_X_Mangoum permethrin alive versus dead. The Fst values were calculated within a window of 100kb and 50kb window steps using winScan and Fst per SNP generated in popoolation2. (C) Manhattan plot showing Cochran-Mantel-Haenszel (CMH) test or genome wide association analysis between F4 Kisumu_X_Mangoum permethrin alive and F4 Kisumu_X_Mangoum permethrin dead taking into account all the replicates. This test detected consistency in allele frequency changes in biological replicates and validated that signals in chromosomes 2L, 2R were the main drivers of permethrin resistance in our data set. (D) Neutrality test (Tajima’s D) within the 300kb CYP6P3 cluster computed within 5kb window and 2.5kb step. (E) & (F) Nucleotide diversity (pi) along the 300kb and 10kb CYP6P3 clusters respectively computed within a 5kb window and 2.5kb step and 500bp window and 250bp step.

Copy number variations (CNVs) in the 2R chromosome were predicted using the hidden Markov model (HMM) with normalized coverage across the chromosome. No CNV was found within or around the CYP6 cluster. However, regions with copy number greater than 3 were detected across Chromosome 2R (Fig. S5 **A**). Visualization of these regions with the Samplot package (https://github.com/ryanlayer/samplot) confirmed duplications around the *ace*1, beta-glucuronyltransferase (AGAP001367), Protease m1 zinc metalloprotease (AGAP013155) genes showing relatively higher coverage in permethrin alive (resistant) compared to susceptible dead and the Kisumu (Fig. S5 **B**, **C**, **D**). We also identified some inversions in the putative allatostatin receptor 2 (GPRALS2) (Fig S5. **e**) though not associated with resistance. Indels were also called using Manta (Chen et al., 2016) in the 2R chromosome and Fst was used to check their association with the resistant phenotype. Although this analysis revealed outlier (95-99% quantiles) Fst values ranging from 0.6-1.0 (Fig. S6 **A**), samplot visualization showed only one indel with resistance association. A 473 bp insertion in the zinc finger protein GLIS1/3 (AGAP003885) gene was found exclusively in resistant mosquitoes (Fig. S6 **A**). All the other indels showed no link to resistance (Fig. S6 **B**) and were mostly found in non-coding regions spanning uncharacterized genes (unspecified products).

Given that no structural variations (SVs) were found within or close to the CYP6 cluster, we subsequently focused on the non-synonymous mutations that might be driving resistance in this locus. A total of 377 non-synonymous mutations were detected among which 68 (18.03%) were associated with resistance as they were found to have higher frequency in resistant populations compared to susceptible populations (Fig 2). Those mutations were mostly found in cytochrome P450 genes (*CYP6AA1*, *CYP6AA2*, *CYP6P15*, *CYP6P1*, *CYP6P2*, *CYP6P3* and *CYP6AD1*) in addition to the carboxylesterase *COEEAE60*, zinc finger-containing protein and cathepsin-F genes. The *kdr* mutation L1014/995F was fixed in the field mosquitoes. Frequencies of 56.9%, 7.0% and 0.0% were recorded in resistant hybrids, susceptible hybrids (P=0.0011, Fisher exact test) and Kisumu (P<0.001, Fisher exact test) respectively confirming moreover the re-segregation of genotypes in the hybrid population (Fig. 2). Missense mutations with association with resistance phenotype were also identified in the GSTE cluster with the main one (Lys190Glu) found in *GSTE4* with the 190Glu frequency as high as 44% in resistant hybrid mosquitoes compared to 10% and 0.0% in susceptible hybrids (P = 0.02, Fisher exact test) and Kisumu (P = 0.0014, Fisher exact test), respectively. Due to its high content of non-synonymous mutations and signs of selection, more attention was paid to the CYP6 cluster with 10 genes (7 P450s) showing significant allelic variation between alive and dead F_4_ mosquitoes. This high number of associations is likely a result of the linkage or hitchhiking between the main variants and neighboring ones. To ascertain which gene was the most likely driver of this metabolic resistance, we hypothesized that it must be a P450 due to synergist recovery with PBO in this location and the abundance of CYP6 at this locus. As Poolseq data might not always provide a very precise allelic frequency due to biases associated with pooling of mosquitoes, we opted to look at historical reports and orthologs to other species. From this approach, the *CYP6P3* was chosen as the most likely candidate resistance gene. Indeed, *CYP6P3* is the main gene out of the 7 associated P450s in this locus to have been shown to be able to metabolise pyrethroids (Muller et al 2008, Edi et al 2014) highlighting the likelihood that polymorphisms in this gene could be most likely drivers of resistance to pyrethroids. *CYP6P3* also presented a single non-synonymous mutation E202D with association with insecticide resistance (Fig 2, Data_file-sheet01). Its frequencies were 87.50%, 70.83%, 35.80%, 4.35% and 6.59% in Mangoum 2022, Mangoum 2018, resistant hybrid, susceptible hybrid and Kisumu mosquito populations respectively (P<0.001). The increase in the frequency of this mutation between 2018 and 2022 suggests continuous selection in the field (*21, 22*).

**Figure 2:**
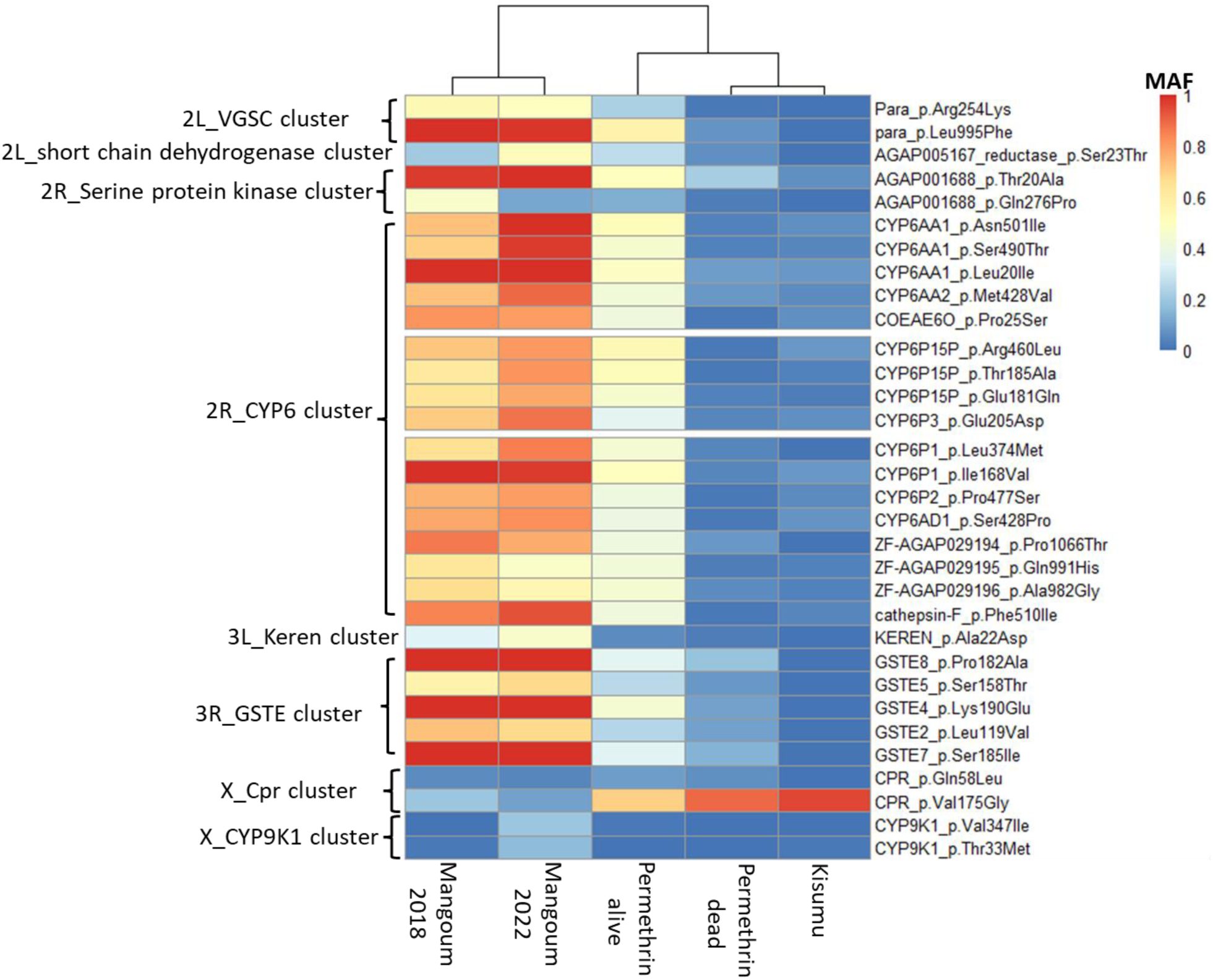
Minor allele frequency of key non-synonymous mutations located in genes within the different clusters of interest between study populations. Vcf file was generated using varscan2 and the annotation and extraction of non-synonymous mutations were carried out using the SnpEffect package.

### Comparative analysis of *CYP6P3* polymorphism between Mangoum and Kisumu

To further establish the association between *CYP6P3*-E205D substitution and resistance phenotype, we cloned and sequenced its full coding region using cDNA from Mangoum and Kisumu. The Mangoum *CYP6P3* exhibits a lower genetic diversity with a predominant haplotype containing the E205D mutation (Fig S7) contrary to Kisumu which exhibited a high polymorphism with no predominant haplotype. The *CYP6P3* polymorphism was further assessed by Sanger sequencing 2500bp genomic DNA fragments spanning the 5’UTR (1kb) + *CYP6P3* (full-length). Fifty DNA samples (20 hybrid permethrin resistant, 18 hybrid permethrin susceptible, and 12 Kisumu controls) were successfully sequenced. The nucleotide diversity in the coding region was higher in Kisumu and susceptible hybrids, at 0.0097 and 0.01 respectively, compared to the resistant ones (0.0053). Haplotype diversity was also lower in alive mosquitoes (0.68), compared with 0.99 and 0.93 in Kisumu and susceptible hybrid mosquitoes respectively, suggesting that allelic variation at this gene is associated with pyrethroid resistance. The same trend was observed for the number of polymorphic sites and haplotype number with the alive group having the lowest (19 and 11 respectively) compared to 28 and 18 in dead hybrid mosquitoes (Table S3). The same pattern was observed when the intronic region only or the coding and intronic regions were analyzed (Table S3).

A total of 47 haplotypes were observed across the whole population for both the coding and coding + intronic regions resulting in 16 different protein sequences, of which 8 were unique to Kisumu (Data_file-sheet02). Of the 47 haplotypes, 11 were found in resistant mosquitoes, with 4 protein sequences, of which the sequences with G615C polymorphism (205D) mutation are the predominant (29/40), (Fig. 3 **E**, Data_file-sheet02). This mutation was completely absent in the remaining 12 protein variants found predominantly in Kisumu and susceptible hybrid mosquitoes indicating that the 205D mutation is significantly associated with pyrethroid resistance. This association was further portrayed by the phylogenetic and haplotype network analyses with the clustering of alive mosquitoes in the tree and the presence of a major haplotype (H3) composed essentially of alive mosquitoes (Fig. 3 **A** & **C**). Moreover, the resistant haplotypes were more homogeneous, clustering around the main haplotype H3 (less mutational step differences ≤7). Kisumu was the most diverse population with 21/47 haplotypes composed mainly of singletons (18/21). Maximum likelihood tree and haplotype network of *CYP6P3* intronic region established similar clustering with a major resistant haplotype harboring the 1171C mutation (Fig. 3 **B** & **D**). Analysis of the coding and intronic region also revealed a predominant resistant haplotype (28/40) carrying both 615C (205D) and 1171C mutations (Fig S8.) suggesting the two mutations are linked.

**Figure 3:**
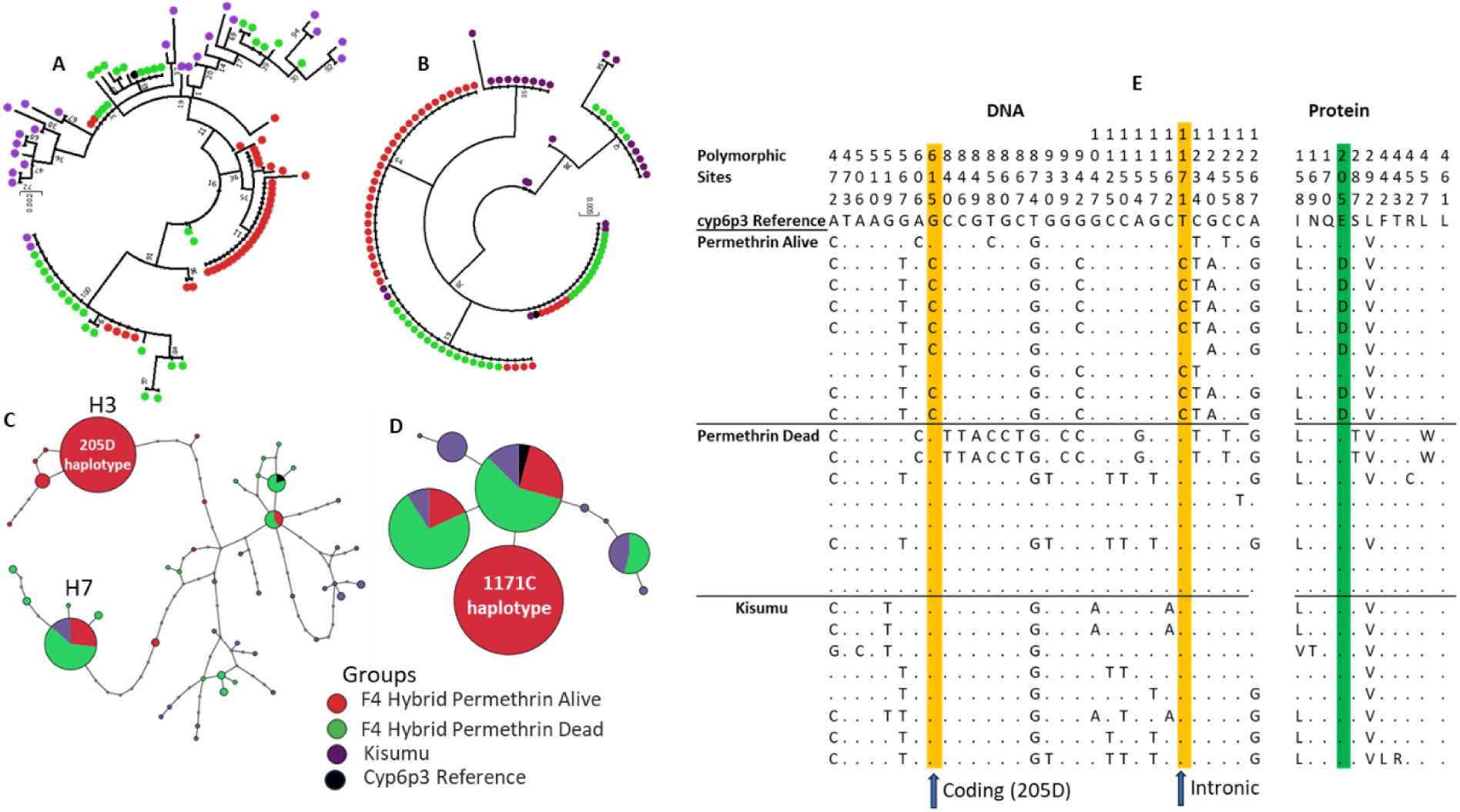
CYP6P3polymorphisms: (A) Maximum likelihood tree of CYP6P3 coding region with F4 Kisumu_X_Mangoum hybrid mosquitoes exposed to permethrin alive (resistant) and dead (susceptible) as well as Kisumu. (B) Same as in (A), but with the intronic region. (C) CYP6P3 coding region haplotype network (TCS) between Kisumu, susceptible and resistant mosquitoes. (D) Same as in (C), but with the intronic region. The pie size reflects the haplotype frequency. Each node represents a mutation. (E) MEGA-aligned CYP6P3 DNA and protein sequences from resistant, susceptible and Kisumu mosquitoes showing the different polymorphisms. Polymorphic sites highlighted in yellow in genomic sequences represent the G615C and T1171C in the introns with high frequency in resistant mosquitoes. The G615C causes a nonSyn mutation E205D highlighted in green in protein sequences. The tree was built using MEGA v7.0 with 1000 bootstraps.

### Prediction of the impact of the *CYP6P3*-E205D mutation using modelling and docking

Docking simulations were performed to determine if the *CYP6P3*-E-205D allelic variation could impact the binding capacity and/or interactions of *CYP6P3* with pyrethroid insecticides. These simulations revealed that both *CYP6P3*-E205 and *CYP6P3*-205D models bind to α-cypermethrin and permethrin with low free energy of binding and high affinity (Table S4) suggesting the abilities to metabolize pyrethroids (though with α-cypermethrin exhibiting the highest score of -11.2 kcal/mol in the active site of *CYP6P3*-205D model).

Permethrin docked into the active site of *CYP6P3*-E205 with phenoxy oxygen above the heme and with the C-2 of the benzyl group at 6.3Å from the heme, while C-4′ of the phenoxy ring is facing away from the heme at a distance of 8.5Å (Fig. S9 **A**). Energetic contributions include hydrophobic interactions, contributed by Phe^123^ (of the substrate recognition site 1, SRS-1), Val^310^ and Leu^313^ of SRS-2, Val^380^ (SRS-5) and Ile^494^ (SRS-6); hydrogen bond by Glu^215^ (SRS-2) and a pi-stacking between Phe^123^ (SRS-1) and the phenoxy- and the benzyl rings. Permethrin docked into the active site of *CYP6P3*-205D with the ester oxygen above the heme suggesting ester hydrolysis. The C2 of benzyl group is located 8.5Å from the heme iron and while C-4′ of the phenoxy ring at 11.6Å (Fig. S9 **B**). In this pose no residue was shown to be involved in hydrogen bonding, instead hydrophobic interactions were predicted primarily by network of aromatic rings, including His^107^, Phe110 and Phe^123^ all from SRS-1, in addition to Glu^215^ (SRS-2), Leu^313^ (SRS-4), Thr^318^ (SRS-4) and Val^380^ (SRS-5). As above Phe^123^ was involved in pi-stacking with both the phenoxy and the benzyl rings.

In *CYP6P3*-E205 model α-cypermethrin binds with the carboxyl ester group oriented above the heme and the gem trans methyl group of cyclopropane ring at 5.6Å from the heme iron (Fig. 4 **A**, Fig. S9 **C**). In this posture the C4′ is located at 12.4Å from the heme iron, suggesting that first hydroxylation is not on the aromatic rings. Energetics were contributed through hydrophobic interactions involving Ile^59^, Phe^123^ of SRS-1 (alone exhibiting 4 interactions), Glu^215^ (SRS-2), Ile^313^ and Thr^318^ (both from SRS-4), Pro^379^, Val^380^ and Glu^381^ (all three residues from SRS-5), and Thr^495^ (SRS-6). Two residues, Glu^215^ (SRS-2) and Thr^318^ were involved in hydrogen bonding with this insecticide. Similar pattern was observed in the *CYP6P3*-205D model, α-cypermethrin binding with the gem dimethyl group of the cyclopropane ring oriented above the heme, with the trans methyl group at 5.3Å from the heme iron (Fig. 4 **B**, Fig. S9 **D**). In this posture the C4′ is located at 10.8Å from the heme iron, suggesting that first hydroxylation is not on the aromatic rings. Energetics were contributed only through hydrophobic interactions involving Phe^123^ of SRS-1 (alone exhibiting 5 interactions), Glu^215^ (3 interactions) and Leu^216^ both from SRS-2, Ala^314^, Thr^321^ and Thr^318^ (all three residues from SRS-4), Pro^379^ (SRS-5) and Ile^494^ and Ile^496^ (both from SRS-6).

**Figure 4:**
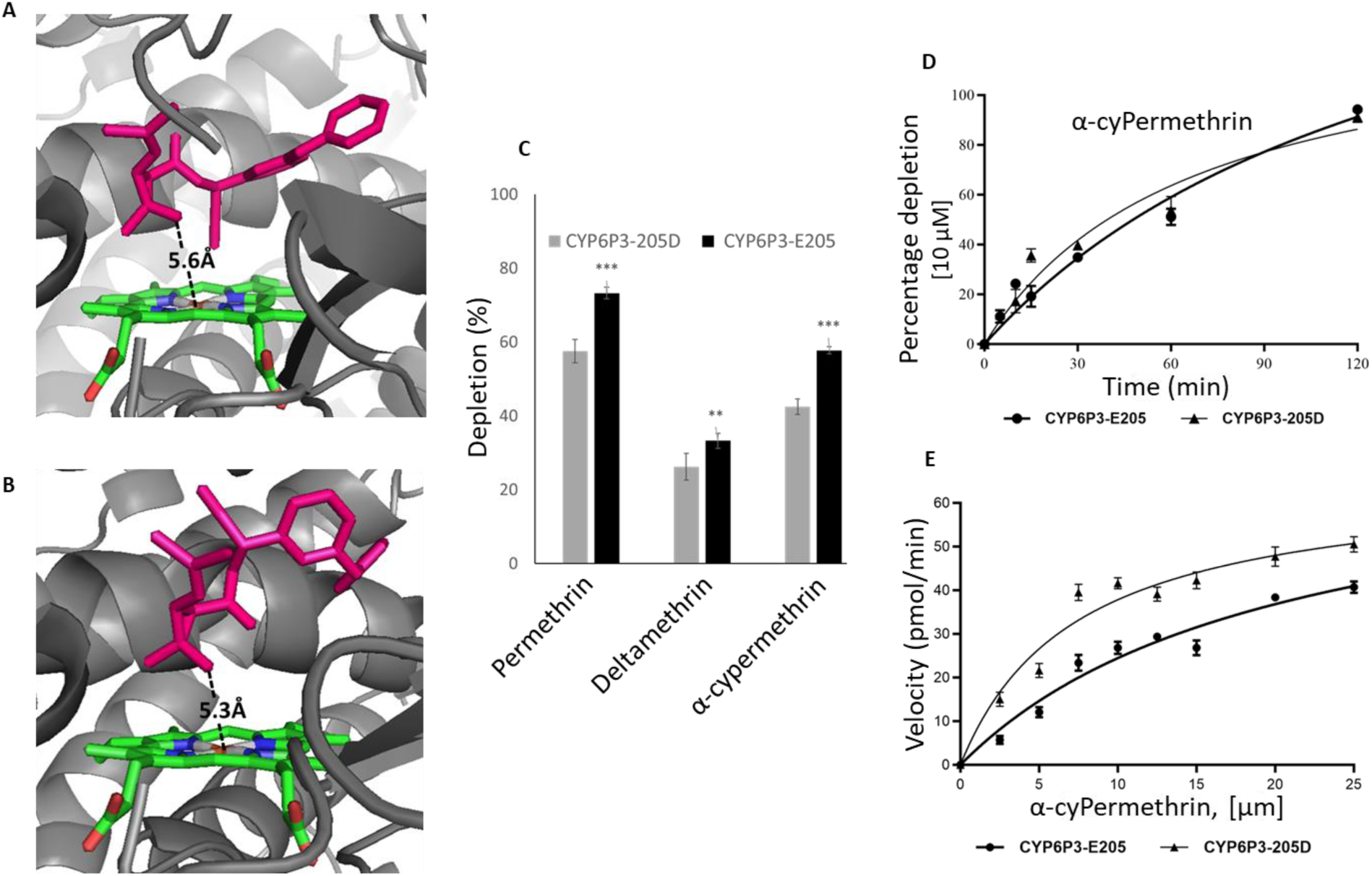
In silico and in vitro functional validation. Predicted binding mode of (A) α-cypermethrin in CYP6P3-E205, (B) α-cypermethrin in CYP6P3-205D models. CYP6P3 helices are presented in grayscale 50, heme atoms are in stick format and coloured by atoms, α-cypermethrin in hot pink. Distance between possible sites of metabolism and heme iron are annotated in Angstrom. (C) Percentage depletions of type I and II pyrethroid insecticides, results are presented as mean±STD of three replicates each in the negative (-NADPH) compared with the positive (+NADPH). (D) Michaelis-Menten plot for the turnover for α-cypermethrin. (E) Steady-state kinetic parameters for the metabolism of α-cypermethrin by CYP6P3 alleles.

### *In vitro* assessment of the impact of the *CYP6P3*-E205D allelic variation on the ability to metabolize pyrethroids

#### Recombinant CYP6P3 alleles metabolism of insecticides

Mutagenesis and heterologous protein expression were carried out to compare the metabolic profile of CYP6P3-E205 and that of CYP6P3-205D proteins. Initial metabolism assays revealed that both proteins can metabolize pyrethroid insecticides, with higher permethrin depletion observed for CYP6P3-E205 (73.2%±1.6), compared with the substrate that disappeared from incubation with CYP6P3-205D (57.5%±3.1) (Fig. 4 **C**). α-cypermethrin was also metabolised by both CYP6P3-205D and CYP6P3-E205 alleles depleting 42.4±2.1 and 57.7%±1 of this type II pyrethroid, respectively. Both alleles showed a moderately lower depletion of deltamethrin with 26.3%±3.6 for CYP6P3-205D and 33.2%±2 for the CYP6P3-E205 allele (Fig. 4 **C**).

#### Determination of steady state kinetic parameters

The metabolism of pyrethroids followed the Michaelis-Menten fashion with the Mangoum recombinant protein (CYP6P3-205D) exhibiting slightly higher turnover (2.69 min^-1^ ± 0.75) for α-cypermethrin, which is ∼1.2 times higher than the turnover (2.26 min^-1^ ± 0.63) obtained from the CYP6P3-E205 protein (Fig. 4 **D**). For permethrin, the 205D protein also demonstrated faster reaction rate with a turnover value of 2.28 min^-1^ ± 0.90 which is 1.3 times higher than the 1.74 min^-1^ ± 0.14 value obtained with the E-205 allele (Fig. S9 **E**).

Contrasting patterns were also observed for the catalytic constant and affinities. For example, while the E-205 allele exhibited a slightly higher K_cat_ of 1.65 min^-1^ ± 0.32 for α-cypermethrin, compared to the 205D allele (K_cat_ = 1.46 min^-1^ ± 0.15), the 205D allele exhibited three times higher affinity (K_m_ = 7.35 µM ± 2.01) compared with the E205 allele (K_m_ = 20.42 µM ± 7.03). This leads to a higher α-cypermethrin catalytic efficiency (K_cat_/K_m_ = 0.20 min^-1^ μM^-1^ ± 0.06) for 205D allele against 0.081 min^-1^ μM^-1^ ± 0.032 observed in the E-205 allele (Fig. 4 **E**), a ∼2.5 times difference. In the case of metabolism of permethrin (Fig. S9 **F**), the two proteins demonstrated comparable catalytic rates with K_cat_ values of = 1.58 min^-1^ ± 0.13 and 1.57 min^-1^ ± 0.57 for the 205D and E-205, respectively. Just like in the case of α-cypermethrin, the 205D exhibited almost four-times more affinity towards permethrin with a K_m_ value of 7.10 µM ± 2.23 as compared to the 28.02 µM ± 6.05 exhibited by the E-205 allele. This leads to ∼3.7 times higher permethrin catalytic efficiency in the 205D allele (0.22 min^-1^ μM^-1^ ± 0.07) compared to the E-205 allele (0.06 min^-1^ μM^-1^ ± 0.02).

### *CYP6P3*-205D allele confers resistance to different classes of insecticides in transgenic Drosophila flies

To find out whether the expression of the resistant allele of *CYP6P3* can contribute to insecticide resistance, *D. melanogaster* flies expressing the Mangoum *CYP6P3*-205D allele were generated and exposed to different classes of insecticides. Assessing resistance level of the transgenic line revealed lower mortality rates with flies expressing the *CYP6P3*-205D compared to the control line for both type I and type II pyrethroids (Fig. **5 A**, **B** & **C**). Flies expressing the *CYP6P3*-205D allele were significantly more resistant to permethrin with significantly lower mortalities than control flies at different time-points post-exposure and a final mortality of 39.13% ± 4.8 after 24 h compared to 83.73 % ± 3.6in the control group (p = 0.000038) (Fig. 5 **A**). Resistance was also observed against the type II pyrethroids with the *CYP6P3*-205D flies exhibiting significantly lower α-cypermethrin mortality rates at different time-points, and a final mortality of 56.23% ± 2.88 after 24 h, compared with 87.56% ± 3.22 in the control group (P < 0.0001). Similar pattern was observed for deltamethrin with significantly lower mortality rates at 3h, 6h, 12h and a final mortality of 78.37% ± 5.78 at 24 h compared with 93.03% ± 1.35 (P < 0.0001) in the control (Fig. 5 **B** & **C**).

**Figure 5:**
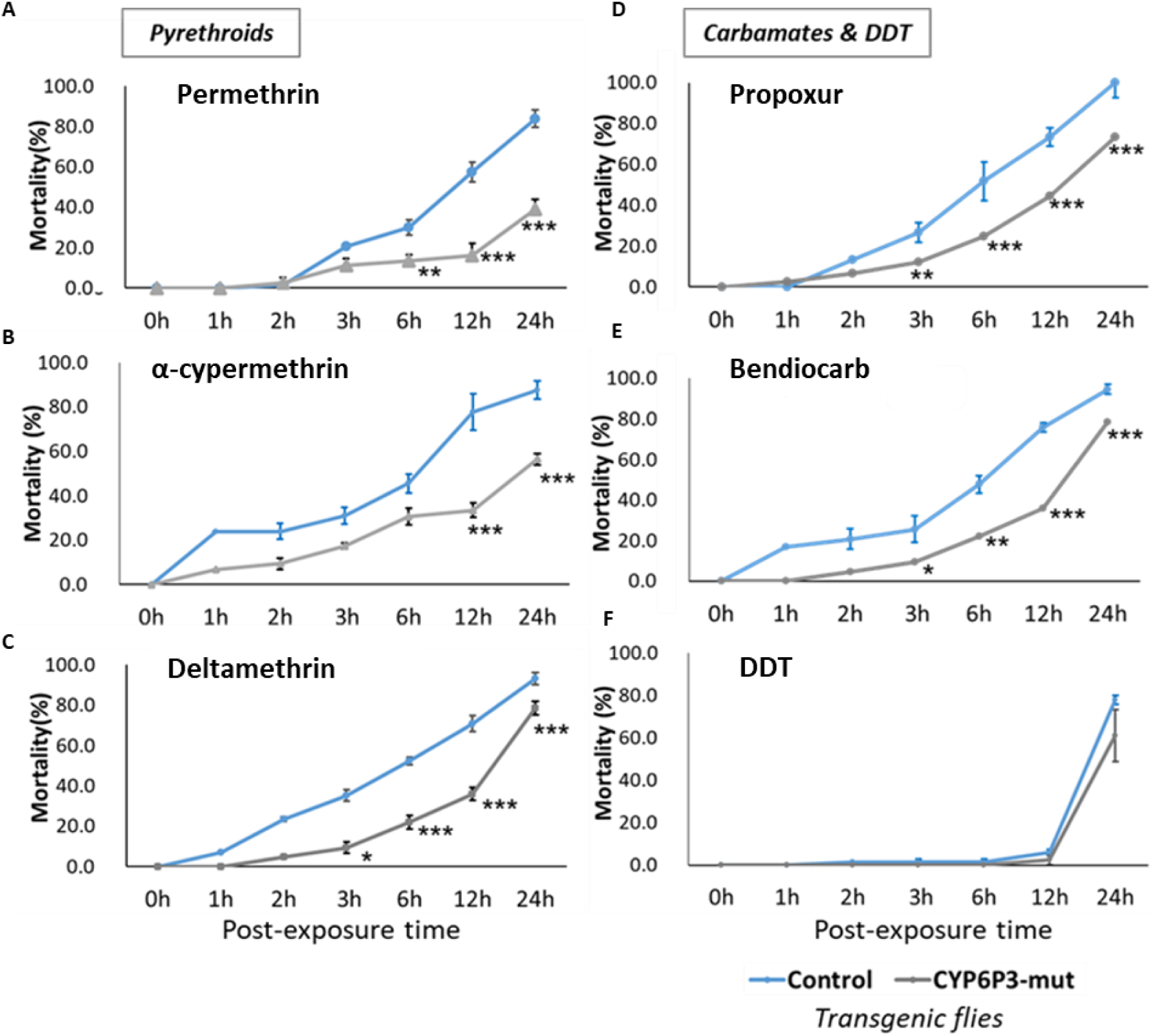
Susceptibility profile of CYP6P3 mut expressing transgenic flies exposed to different insecticide classes. The flies were exposed to a) 4% permethrin, b) 0.007% α-cypermethrin, c) 2% deltamethrin, d) 0.1 % propoxur, e) 0.07% bendiocarb, f) 4% DTT.

Other non-pyrethroid insecticides were also tested. Exposure to the carbamate insecticides, 0.1% propoxur and 0.07% bendiocarb led to mortality rates of 73.34% ± 2.56 and 78.37% ± 1.57 respectively, for *CYP6P3*-205D flies compared to 94-100% mortalities obtained in the control group (P < 0.001, Fig. 5 **D** & **E**). However, no significant differences in mortalities were observed between *CYP6P3*-205D flies and control flies when exposed to DDT, suggesting that this gene does not confer resistance to this organochloride insecticide (Fig. 5 **F**).

### *CYP6P3*-based diagnostic tools design

The observed association of the *CYP6P3-E205D*(G615C) mutation with pyrethroid resistance from genomic analyses and in functional *in vitro* and *in vivo* validation led us to design diagnostic tools to track its spread in the field. An LNA probe-based Taqman assay was designed (Fig. S10 **A**) and was able to successfully differentiate the genotypes (homozygote resistant allele CC (RR), heterozygote CG (RS), and homozygote susceptible GG (SS)) as shown on figure 6**A**. The mutation T1171C in the intron also created a restriction site for the enzyme *MspA1I*. A RLFP-PCR assay was designed (Fig. S10 **B**) and shown to distinguish between the three genotypes, RR, RS and SS (Fig. 6 **B**). For the RFLP assay, digestion of the 1050 bp *CYP6P3* PCR amplicon yielded 2 bands of 600 bp and 450 bp with samples having the CC (RR) genotypes, with heterozygotes, TC (RS) exhibiting three bands, while the wild type TT (SS) genotypes remained undigested.

**Figure 6:**
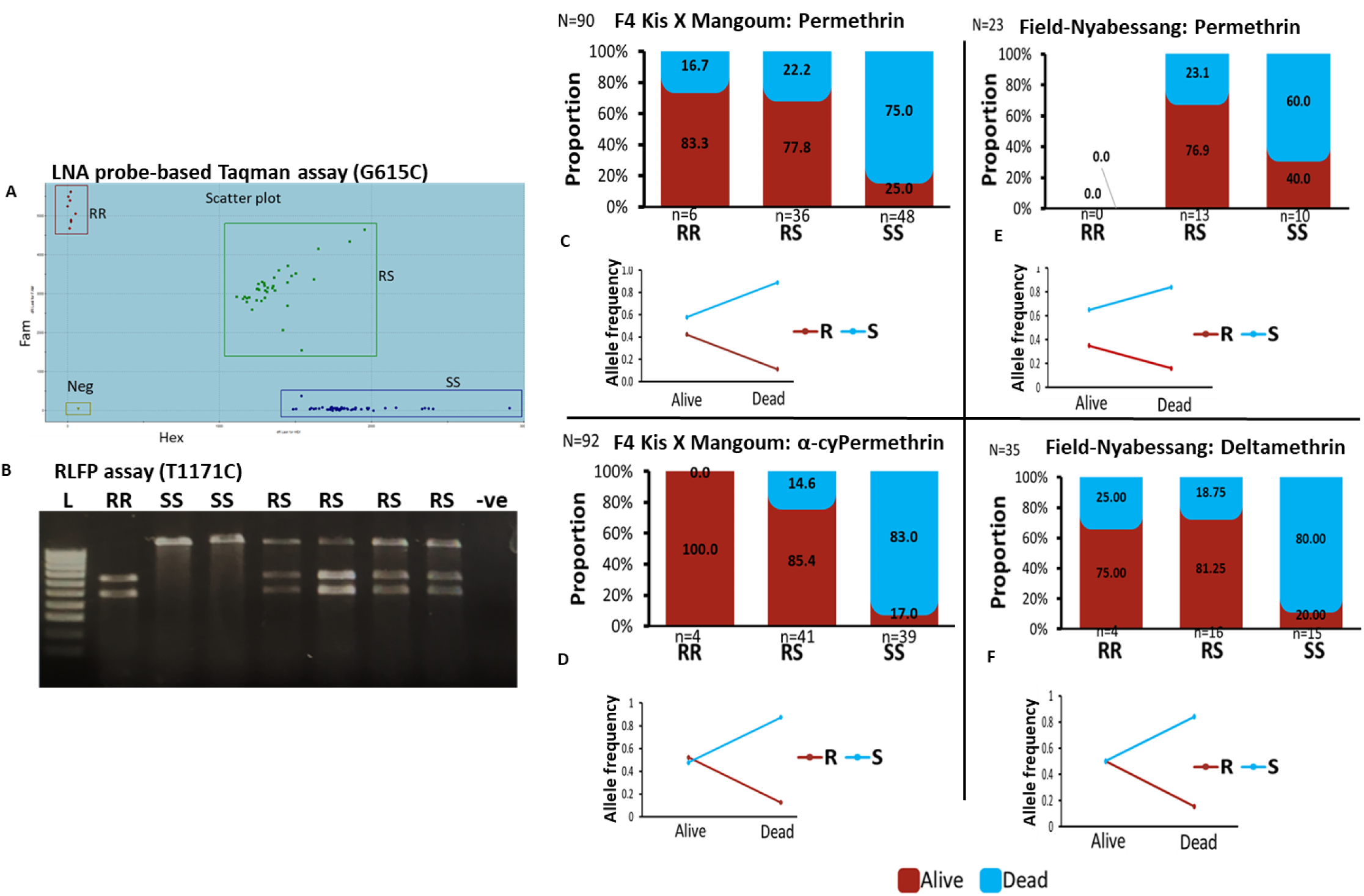
Design and validation of CYP6P3-based diagnostic tools. (**A**) Scatter plot showing clear segregation of genotypes from LNA TaqMan tool for the G615C (E250D) Mutation. This tool was able to delineate the 615C or 205D (resistant, RR) allele from the susceptible 615G or 205E susceptible allele (SS) as well as heterozygotes (RS). (**B**) Gel picture showing the various genotypes following the RFLP assay designed for the CYP6P3 T1171C mutation in the intron, L=Ladder, -ve= negative control. (**C)** & (**D**) Association between the CYP6P3 E205D markers and permethrin and α-cypermethrin respectively using hybrids. (**E)** & (**F**) same as in (c & d) but with field mosquitoes. These assays were carried out using the CYP6P3-205D LNA assays with DNA extracted from F3/4 Kisumu_X_Mangoum hybrid and field (Nyabessang) mosquitoes exposed to different insecticides, Kis = Kisumu.

### *CYP6P3*-based diagnostic tools delineate pyrethroid-resistant mosquitoes from susceptible ones

To confirm the association between the 205D mutation and the resistance phenotype, permethrin-, α-cypermethrin-, and deltamethrin-exposed hybrid mosquitoes (alive and dead) were genotyped using the LNA-based tool. For permethrin, the resistant allele (R) was more frequent in alive (0.48) than in the dead (0.17) mosquitoes (Fig. 6 **C**) with the R allele conferring more chances to survive than the S (OR: 5.9, CI: 2.8-12.3, p<0.0001). About 83.3% (OR:15, CI: 1.6-141.5, P = 0.0180) and 77.8% (OR: 10.5, CI: 3.8-29.2, p<0.0001) of mosquitoes with the RR and RS genotypes respectively, were alive compared to 25% in the SS genotype group (Fig. 6 **A**). The same trend was observed with mosquitoes exposed to the type 2 pyrethroid α-cypermethrin and deltamethrin (Fig. 6 **D**; Fig. S11 **A**). Mosquitoes with the RR genotype were 41 and 43 times more likely to survive than those with the SS genotype for α-cypermethrin (OR: 41.8, CI: 2.05-852.1, P = 0.0152) and deltamethrin (OR: 43.4, CI: 4.3-432.2, P = 0.0013), respectively.

### 205D is strongly associated with pyrethroid resistance in the field in Cameroon

Genotyping of pyrethroid-exposed field collected mosquitoes in Cameroon (Nyabessang) with the LNA assay produced similar pattern (Fig. 6 **E** & **F;** Fig. S11 **B**) with mosquitoes bearing the RR and RS genotypes surviving better than their SS counterparts. However, significant results were obtained with deltamethrin only probably due to small sample size of field samples. For deltamethrin, 75% (OR: 12, P0.06) and 81.25% (OR:17.33, p=0.0017) of RR and RS were alive compared to 20% in the SS group. The same profile was seen with permethrin and α-cypermethrin (Fig. 6 **E**; Fig. S11 **B**).

Genotyping the T1171C marker in permethrin-exposed hybrid mosquitoes using the RFLP assay also showed a clear association of the marker and pyrethroid resistance Fig. S11 **C**). The resistant allele C (R) was more frequent in alive (0.45) than in dead (0.05) mosquitoes (OR:15.8, CI:5.9-42.3, P < 0.0001). To determine whether there was an agreement between the two diagnostic assays developed, a set of 29 samples was genotyped using the *CYP6P3-E205D*LNA and T1171C-*CYP6P3* RFLP assays. The assays were found concordant (R^2^=0.95, P = 6.375e-14, Fig. S11 **D**).

### Combined effect of *CYP6P3* and *kdr* on pyrethroid resistance

As the two major genomic signals were located across the sodium channel gene (kdr; 2L) and the CYP6 cluster (CYP6P3; 2R), we opted to assess whether CYP6P3 was combining with kdr to confer greater level of resistance to pyrethroids, both markers were genotyped in pyrethroid-exposed hybrid mosquitoes (92 dead and 92 alive). An analysis of the contribution of each marker to pyrethroid was first performed revealing that each marker was associated with pyrethroid resistance (Fig. S12 **A**-**F**). The CYP6P3-205D allele confers more than two-fold a greater level of resistance to permethrin (OR:26.4, P = 0.0023, Fig. S12 **A**) than kdr (OR:12.1, P = 0.0001) when comparing the RR genotypes to the SS for both markers. However, a different trend was found with type 2 pyrethroids. The CYP6P3-205DRR genotype had reduced ability to confer resistance to α-cypermethrin (OR:57.0, CI: 3.2-1015.8, P = 0.0059) and deltamethrin (OR:43.4, CI: 4.3-432.2, P = 0.0013) when compared to the 1014F-kdr RR genotype which gave more chances to mosquitoes to survive α-cypermethrin (OR:64.5, CI: 13.2-315.3, P < 0.0001, Fig. S**9**) and more so for deltamethrin exposure (OR: 406.7, CI: 19.9-8282.6, P = 0.0001, Fig. S12**c**, Table S5).

We next assessed the combined impact of the two markers (*CYP6P3-205D & kdr-1014F)* in our study population revealing that both markers combine additively to confer higher level of resistance. No RR/RR was observed for permethrin and α-cypermethrin and as such the RR/RR genotype was not considered in the analysis. For permethrin, Mosquitoes with the RR/RS genotype (homozygote resistant for *CYP6P3* and heterozygote for *kdr* or vice versa) had 64.8 times more chances to survive permethrin exposure compared to those with the double susceptible (SS/SS) genotype (OR: 64.8, P<0.0001, Fig. 7 **A**). Interestingly, the ability to survive permethrin exposure increased by 4-5 times in mosquitoes heterozygote (RS/RS) for both markers in comparison of being heterozygote for only one and susceptible to the other (RS/SS or SS/RS) (OR: 51.8, P < 0.0001) implying an additive effect between the two resistance mechanisms (Fig. 7 **A**). Mosquitoes homozygote for only one marker and susceptible to the other (RR/SS genotypes) had also greater chances (OR: 45, P < 0.0008, Table S5) to survive permethrin exposure compared to double SS/SS genotypes. However, the sample size for the RR/SS genotype was small and this may have affected the results. Similar observations were made for α-cypermethrin-and deltamethrin-exposed mosquitoes (Fig. 7 **B**, Fig S13 **A**, Table S5). A similar assessment was done with the field samples from Nyabessang where the *kdr* is nearly fixed (77/80). We noticed that homozygote RR mosquitoes for *kdr* and heterozygote for *CYP6P3* (RS/RR) were more likely to survive deltamethrin exposed compared to those with *kdr* RR genotypes (SS/RR) only (OR:10.1, P=0.008, Fig. 7 **C**). The same profiles were seen with mosquitoes exposed to permethrin and α-cypermethrin but the association was not significant due to small samples size (Fig. S13 **B** & **C**). It was difficult to have dead mosquitoes and SS/SS genotypes in the field because of the strength of the resistance.

**Figure 7:**
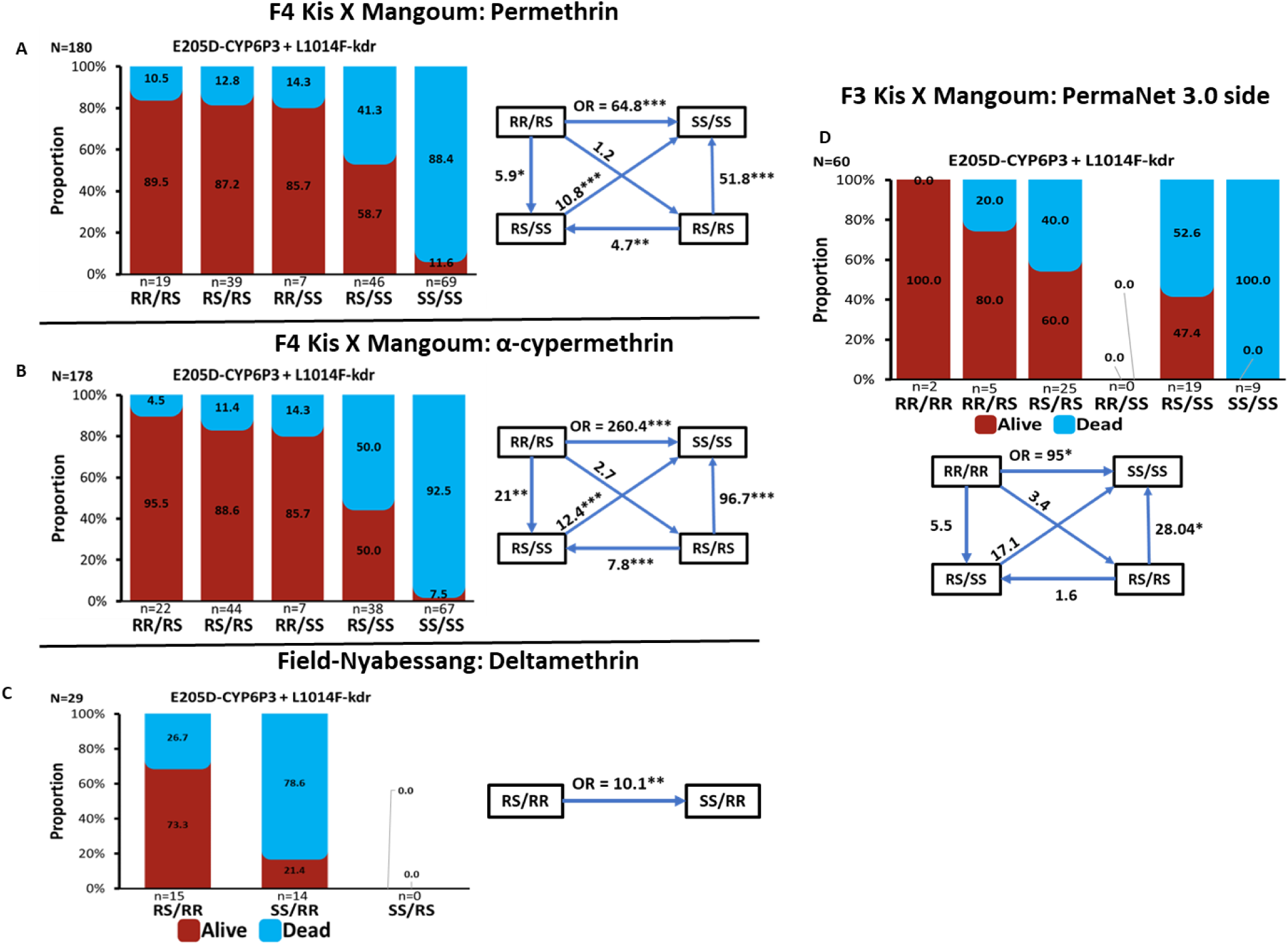
Impact of CYP6P3-E205D and kdr-L1014F markers on pyrethroid insecticide resistance and efficacy of control tools. The figure depicts the frequency distribution of CYP6P3-E205D and kdr-L1014F genotypes between dead and alive F3/4 hybrid *An. gambiae* after exposure to 1x permethrin (**A**), 1x α-cypermethrin (**B**) using WHO bioassay. **C**) same as in (**A** & **B**) but with field mosquitoes exposed to 1x deltamethrin. **D**) same as in (**A** & **B**) but with F3 hybrid *An. gambiae* exposed to PermaNet 3.0 side using cone assay. It also gives the odds ratio (OR) and associated significance between different genotypes and the ability to survive exposure to the various insecticides and PermaNet 3.0 side. Statistics: “*” = P & 0.05, “**” = P& 0.01, “***” = P &0.001.

### Impact of *CYP6P3* and *kdr* on LLINs using cone assays

To assess the association of *CYP6P3* and *kdr* on the efficacy of LLINs, both markers were genotyped in the same population of hybrid mosquitoes exposed to different insecticide-treated bednets. The LLINs included pyrethroid only nets: Olyset (permethrin), Interceptor (200 mg/m2 α-cypermethrin), DuraNet (270 mg/m2 α-cypermethrin), and pyrethroid plus: the synergist piperonyl butoxide (PBO) nets, including the Olyset+ (permethrin+PBO), PermaNet 3.0 top panel (deltamethrin+PBO). Apart from PermaNet 3.0 side, all PBO nets recorded 100% mortality and genotyping of *CYP6P3*-205D and *kdr*-L1014F markers revealed the presence of the genotypes (RR, RS and SS) in the dead mosquitoes. The *CYP6P3*-205-D marker was found to be associated with reduced efficacy of the other LLINs tested with more mosquitoes with the RR and RS surviving upon exposure (Fig. S14 **A**, **C** & **E**). The R allele was significantly associated with the ability of mosquitoes to survive exposure to Olyset (OR:2.05, P = 0.0145), Interceptor (OR: 1.98, P = 0.0185) and PermaNet 3.0 side (OR:2.6 P = 0.0015) compared to the susceptible allele (Fig S14, Table S6). The RR genotypes showed increased survivorship for PermaNet 3.0 side compared to their SS counterparts (OR:13.7, P = 0.025). The same trend was observed for Interceptor with the RS genotypes (OR:8.5, P = 0.009). Although the RR also had the ability to survive exposure to Olyset (OR:5.4, P = 0.08) and Interceptor (OR:6, P= 0.1090), this observation was not significant (Table S6).

Having the *kdr*-1014F also correlated with the ability to survive exposure to the LLINs with the correlation being more pronounced with Interceptor and PermaNet 3.0 side. (Fig. S14 **B** & **F,** Table S5). The RR (OR:14.2, P=0.007) and RS (OR:8.2, P=0.0016) genotypes had high survivorship for Interceptor compared to susceptible mosquitoes. Similar observation was made for PermaNet 3.0 side (Table S6). Only the RS genotype showed significant association with Olyset (OR:4.3, P=0.01, Table S6).

### Combined impact of *CYP6P3* and *kdr* on LLINs using cone assays

With respect to the combined effect of the two markers (*CYP6P3-205D & kdr-1014F)*, similar genotypes were merged before analysis to account for the small sample size. Indeed, having 4 resistant alleles (RR/RR) significantly increased likelihood to survive exposure (Fig. 7 **D**, Fig. S13 **D** & **E**) to Olyset (OR:13.0, P = 0.05), Interceptor (OR:115, P = 0.02), and PermaNet 3.0 side (OR: 95.0, P = 0.0319) compared to SS/SS mosquitoes. As seen with tube assays, the ability to survive permethrin exposure increased in mosquitoes heterozygote (RS/RS) for both markers in comparison of being heterozygote for only one and susceptible to the other (RS/SS or SS/RS); OR: 7.04 vs 6.06 for Olyset, OR: 53.6 vs 23.0 for Interceptor and OR: 28.04 vs 17.1 for PermaNet 3.0. This furthermore suggests an additive effect between the two resistance mechanisms in reducing the efficacy of pyrethroid-based nets (Fig. 7 **D**, Fig. S13 **D** & **E**), Table S6).

### Geographic distribution of the *CYP6P3* E205D mutation across Africa

Genotyping of *CYP6P3*-E205D revealed its presence in 4 of the 6 countries (Sierra Leone, Ghana, Cameroon, Uganda) tested in the study. The mutation was absent in Malawi and DRC (Fig. 8 **A)**. Uganda had the lowest frequency (0.05) of the resistant allele (10.8% RS and 89.2% SS) indicating its low spread or absence in the East and towards the Southern Africa (Fig. S15 **A**). The highest frequency (0.80) was recorded in Sierra Leone (Kono) with RR and RS genotypes as high as 67.9% and 25% respectively, followed by Cameroon with overall 39.1% RR and 41.1% RS and a frequency of 0.60. It important to mention that many samples from different localities were genotyped in Cameroon and the mean genotype proportions and frequency are presented in figure 8**A** & Fig. S15 **A.** The *CYP6P3* mutation was found at low frequency (0.15) in *An. coluzzii* with 13% and 4.3% RR and RS genotypes in Ghana (Obuasi) suggesting a possible introgression of this allele in this sibling species.

**Figure 8.**
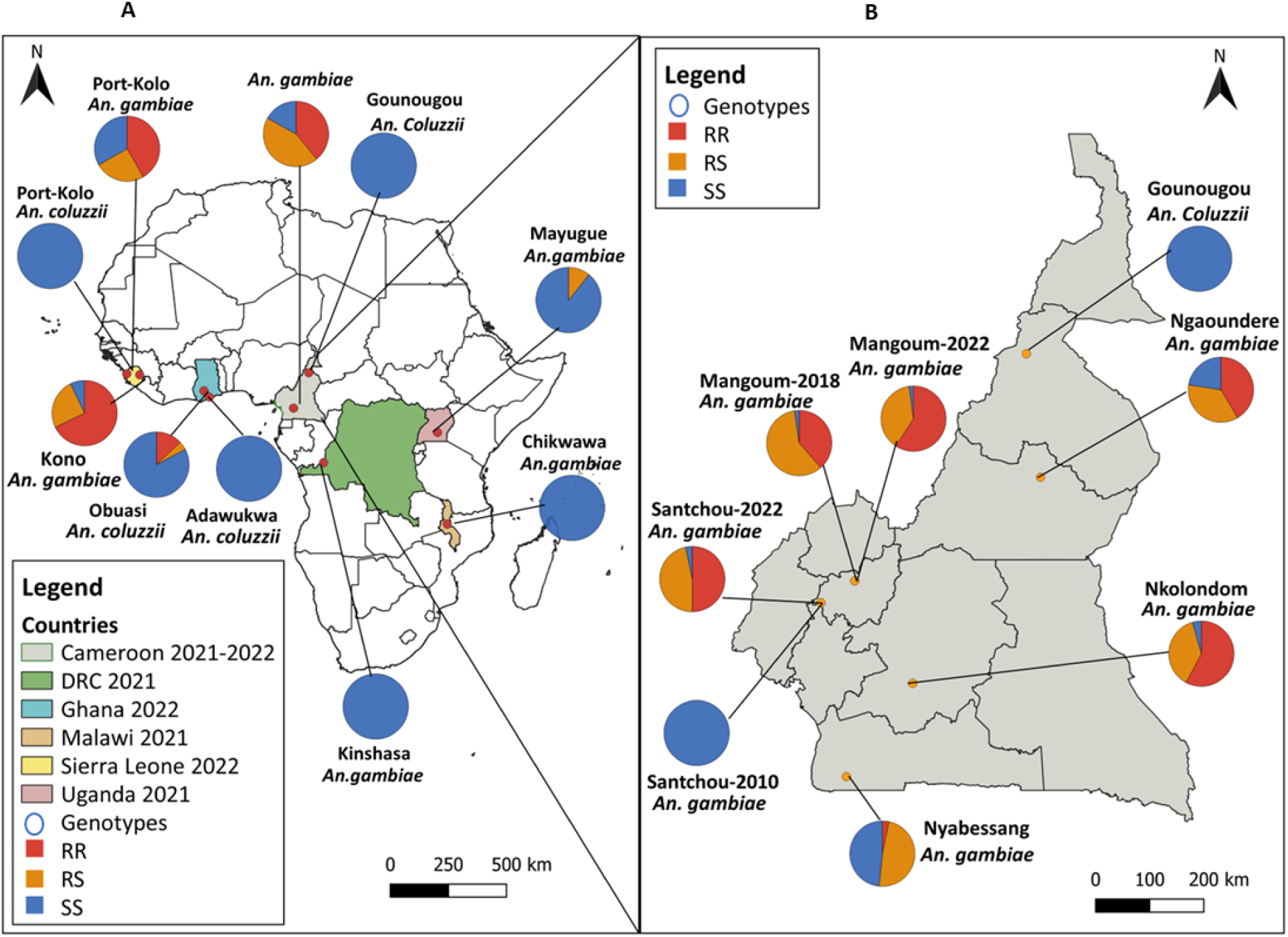
Geographical distribution of the CYP6P3 E205D mutation across Africa. (**A**) Map of Africa showing the genotypic distribution of the CYP6P3 E205D mutation across the continent. (**B**) Map of Cameroon showing the genotypic distribution of the CYP6P3 E205D mutation across different localities. An=*Anopheles*. Maps were generated using the quantum geographic information system (QGIS) v3.14 and open-access shapefiles (https://gadm.org/).

In Cameroon, the mutation was present in all the localities tested except in Gounougou in the northern region where *An. coluzzii* is the main malaria vector species (Fig. 8**B**). The mutation was predominant in Mangoum-2022 (59.18% RR, 38.78% RS and 2.04% SS and Santchou-2022 (50.0% RR, 46.7% RS and 3.3% SS) in the western region with frequencies of 0.79 and 0.73 respectively. Similar proportions were found in Nkolondom (57.8% RR, 37.8% RS and 4.4% SS) in the Centre region with a frequency of 0.77 while Nyabessang (South region) recorded the lowest frequency (0.27) (with the SS genotype as high as 48.8%). Genotyping of samples collected back in 2010 in Santchou revealed a complete absence of the *CYP6P3* E205D mutation suggesting that it must have been selected over years with the continuous implementation of the various vector control measures and use of pesticides for agricultural purposes (*23*) (Fig. S15 **B**). Similar to the poolseq data, an increase in allele frequency from 0.68 to 0.79 was observed between the years 2018 and 2022 in Mangoum. We also noticed a near significant increase of RR from 38% to 58% (P=0.0623, Fisher exact test) and a reduction of RS from 59% to 38% (P= 0.0628, Fisher exact test) suggesting a stronger effect of selection in this population.

## Discussion

Elucidating molecular basis of insecticide resistance is a prerequisite to the implementation of effective insecticide-based control strategies. Lack of DNA-based markers of P450-based metabolic resistance in *An. gambiae* has for long prevented thorough understanding of the dynamic of resistance in this major malaria vector. This study has filled such knowledge gap by identifying the major genetic loci associated with metabolic resistance to pyrethroids in populations from Central/West Africa. Variant E205D in the *CYP6P3* gene within the CYP6 cluster located on chromosome 2R was further shown to be involved in resistance to both type I and II pyrethroids using a combination of techniques. DNA-based diagnostic tools were designed to track the spread of the mutation continent-wide and determine its impact on the efficacy of current vector control tools.

### Major genetic loci associated with both target-site and metabolic resistance mechanisms are the major drivers of permethrin resistance in *An. gambiae*

A good association study requires a case and control groups. It is challenging to meet this condition nowadays because the resistance is fixed in the field settings (*24*) making it difficult to get susceptible mosquitoes from the field. The reference strain Kisumu is often used for that purpose, however its genetic background may be different from that of field mosquitoes making the results suggestive. Crossing of resistant field mosquitoes with the susceptible Kisumu lab strain was used to circumvent the issue in the present study as genotyping of the 1014F-*kdr* showed re-segregation of genotypes in the F_4_ Kisumu_X_Mangoum hybrid mosquitoes. This made the hybrids an ideal population for the poolseq whole genome analysis.

Comparing resistant Mangoum (2018 and 2022) mosquitoes (field) to Kisumu revealed signals in all the chromosomes. Zooming in the regions containing the peaks revealed the presence of genes that have previously been associated with insecticide resistance with the main one being VGSC region in the 2L chromosome. Mutations in the para gene specifically the 1014F-*kdr* is known as the main target site mutation associated with DDT and pyrethroid resistance (*9, 25, 26*). In addition to VGSC region, the pronounced peaks were found within the regions containing genes whose overexpression is known to drive resistance in *Anopheles species* such as the cytochrome P450s (*14, 17, 18*) (CYP6 cluster) (*12, 13*), salivary gland proteins (SG5)(*27*), serine protein kinases in the 2R chromosome, CYP9K1 (*28*) in the X chromosome (*29*), glutathione-s-transferases (GSTE cluster)(*30, 31*) in the 3R chromosome and short chain dehydrogenases (*27*) among others in the 2L chromosome. This suggests that genetic variants close to these genes might be associated with pyrethroid resistance. Interestingly, the same peaks were obtained upon comparing resistant field mosquitoes with the highly susceptible hybrids. This result could be linked to the Kisumu background of the hybrid mosquitoes. However, taking the highly resistant hybrids as comparators significantly reduced or abrogated the peaks confirming that the two groups were resistant but with different intensities. Signals were found only on chromosomes 2L and 2R when hybrids mosquitoes were compared (resistant vs susceptible) indicating that those loci may be the main drivers of pyrethroid resistance. This result supports the major role played by both target-site resistance and metabolic resistance in driving pyrethroid resistance in *An. gambiae* with both peaks having similar intensity (*8, 32*). We went further to assess the genetic diversity in the peak regions. The analysis showed a reduced diversity as well as signs of positive selection in the CYP6 cluster for the field samples as previously reported for P450-based resistance Africa-wide *An. gambiae* samples from the 1000 *An. gambiae* genome project (*33*), in *An. funestus* for CYP6P9a/b genes (*34, 35*), in Drosophila with the CYP6G1 loci (*10, 36, 37*). However, the highly resistant hybrids showed signs of balancing selection from the positive values of Tajima’D (Fig 4). This could emanate from the laboratory conditions and the short propagation period used for hybrids rearing. Analysis of SVs across the 2R chromosome did not detect any with association to resistance around and within the CYP6 cluster using Poolseq. For these reasons, more attention was paid on non-synonymous mutations. The condensation of SNPs within the CYP6 cluster could be suggestive of a hitchhiking phenomenon whereby the presence of many SNPs is driven by a main nearby variant as seen in *An. funestus* for the CYP6P9a/b for which the valley of selective sweep spans around 300kb with reduced diversity throughout the cluster of 15 P450 genes (*34, 35, 38, 39*). A differential frequency of nonsynonymous SNPs was particularly noticeable on *CYP6P3, CYP6AA1* and *CYP6P1* which have all previously been shown to metabolise pyrethroids as well as in the *CYP615P* pseudo P450. However, *CYP6P3* is the gene which has by far been consistently associated with pyrethroid resistance in the field across Central (*13*) and West Africa (*14, 17, 18*). On this rationale, a more thorough focus was put on this gene although further analyses should also be performed on other genes notably *CYP6AA1* which has been also shown to metabolise pyrethroids in *An. gambiae* (*19*) as in *An. funestus* (*40*). The CYP6P4 centred variant from this same P450 cluster recently described in East Africa (*19*) was completely absent in these Central and West African populations of *An. gambiae*. To validate the association of this 2R CYP6 cluster in resistance to pyrethroid, the E205D main polymorphism in *CYP6P3* was selected as a key candidate resistance variant as allelic variation of detoxification genes have previously shown to play key role in resistance to insecticides as seen on GST genes with the L119F-GSTe2 alleles shown to confer resistance to DDT/pyrethroid in *An. funestus* (*41, 42*) and the I114T-GSTE2 conferring similar resistance in *An. gambiae* (*43*). Allelic variation in P450s have also been shown to confer high level of resistance to pyrethroids in other mosquito species such as *An. funestus* with site-directed mutagenesis coupled with *in vitro* heterologous expression and metabolism assays showing that directionally selected CYP6P9a/b alleles containing specific amino acid changes are the main drivers of pyrethroid resistance in southern African populations of *An. funestus* (*21, 44*).

### The E205D mutation enhances *CYP6P3* ability to metabolise pyrethroids and increases pyrethroid resistace

Three different but complementary techniques were used to show the implication of the *CYP6P3*-E205D mutation in insecticide resistance. Transgenic flies overexpressing the mutant allele did not only show resistance to pyrethroids but also to carbamates. *CYP6P3* alongside other CYPs (CYP9K1, CYP6P4 ect) have previously been reported upregulated in pyrethroid and carbamate resistant mosquitoes (*12, 13*). This finding is in line with that of Mugenzi and colleagues who demonstrated the implication of CYP6P9a/b in driving carbamate and cross resistance to pyrethroid in *An. funestus* (*45*). Expression of the mutant *CYP6P3* did not appear to drive DDT resistance. DDT has been shown to be more associated with glutathione s-transferase gene family overexpression (*30, 46*). It is important to point out that, in comparison to the susceptible Kisumu strain, *CYP6P3* is not significantly overexpressed in Mangoum where resistance escalation has been established as well as other P450s on this locus such as CYP6P4 (*23*). This implies that *CYP6P3* contribution to resistance could be more qualitative than quantitative with the Mangoum allele (E205D) driving resistance due to its metabolically more active property. In Indeed, *in silico* molecular modeling and docking which show that the 205D allele exhibits the highest affinity and lowest binding energy compared with E-205 for both type I and II pyrethroids. This finding was in line with our genotyping assays that revealed that RR (205D) genotypes had the highest survivorship than homozygote susceptible mosquitoes (E205) when exposed to α-cypermethrin (OR:57, P=0.005), deltamethrin (OR:43.4, P=0.001) and permethrin (OR:26.4, P=0.002).

Hydrophobic interactions and pi stacking were found to be the most important contributors to bindings in 205D, in contrast to E205 in which hydrogen bonding seem to be important for both α-cypermethrin and permethrin. Hydrophobic interactions are relatively stronger than other weak intermolecular forces (i.e., Van der Waals interactions or Hydrogen bonds) and this might justify the higher affinity and lower binding energy of the *CYP6P3*-205D allele. Interestingly, for all insecticides, particularly α-cypermethrin, the pyrethroids bind with the classical site of metabolism (the phenoxy ring) oriented not facing the heme, rather the cis and trans methyl group of cyclopropane ring seem to be the orientation close to heme. This pathway of metabolism has been shown to be minor compared to the binding with the aromatic rings hydroxylation (*47*). These suggest that either the *CYP6P3* metabolise α-cypermethrin and permethrin inefficiently, or the initial hydroxylation of the methyl group makes the primary metabolite more amenable to further hydroxylation on the aromatic ring. The ester group was also seen oriented over the heme iron suggesting a possibility of ester hydrolysis. In addition to *in silico* analysis, in vitro metabolism assays showed that both E-205 and 205-D alleles were able to metabolise pyrethroids. It is interesting to note that even though higher percentage depletions were obtained for the 2 h reactions with the E-205 allele, faster reaction rates and higher affinities were recorded with the 205D allele supporting *in silico* analysis. This indicates that survival to the insecticides depends majorly on how quickly the enzyme metabolises the insecticides rather than how much it clears over a long period window. This further explain the low expression of CYP6P3 in Mangoum. However, *In vivo* confirmation with transgenic flies would have been ideal but we failed to generate CYP6P3-E205 expressing flies for comparison with their CYP6P3-205D.

*CYP6P3* allele from a resistant population of *An. gambaie* from Ghana (*48*) characterized earlier metabolised pyrethroids including permethrin and deltamethrin just like in the case of these two alleles. However higher turnover values were obtained in this study compared to the previous one except for E-205 allele metabolism of permethrin which appeared to be a little slower. Comparable metabolism of permethrin was observed with CYP6P9b, an orthologue of *CYP6P3* in *An. funestus* but higher depletions of up to 80.9% was recorded for the same P450 in the case of deltamethrin (*21*). Higher metabolism with percentage depletions of more than 90% and 80% for permethrin and deltamethrin were respectively recorded in the case of CYP6P9b (*44*). Furthermore, the metabolism of pyrethroids by the two orthologues in *An. funestus* appeared to proceed faster than the rates observed with the two alleles of *CYP6P3*-205D and *CYP6P3*-E205 (*44*). Putting *in silico* and *in vitro* findings together, it is obvious that the 205D allele is a better metaboliser and must have been selected for that reason. Even if this allele is not overexpressed in our study area, its presence confers a greater ability to resist insecticide. Moyes and colleagues showed that *CYP6P3* and CYP6M2 are better metabolizers of pyrethroids in *An. gambiae* using *in vitro* assays (*12*). So this is likely a case where genomic tools prove better than transcriptomic analyses in detect genetic drivers of insecticide resistance.

### A field applicable DNA-based assay allows tracking the spread of the *CYP6P3-E205D*allele revealing that it reduces bed net efficacy

Simple DNA-based assays that strongly correlate with metabolic resistance have long been challenging to design due to the complexity of this mechanism. Here, we took advantage of the detection of the *CYP6P3*-E205D allele to design a DNA-based assay that now allows tracking of P450-based metabolic resistance in *An. gambiae* notably from West and Central Africa. Individual marker genotyping is ideal for monitoring insecticide resistance (*5, 11*). The *CYP6P3*-E205D-based LNA diagnostic tool designed in this study was very accurate in delineating the three genotypes within hybrid and field mosquito populations. Although RFLP assay was designed to target an intronic mutation (T1171C) belonging to the same haplotype with the E205D mutation, The two assays were greatly comparable and could be used interchangeably. With the relative high cost of Taqman thermocylers, the RFLP assay could be used in settings with limited resources. These markers showed a clear association with type 1 and 2 pyrethroid resistance as shown by high ability of RR and RS genotypes to survive insecticide exposure. Hybrid mosquitoes harbouring the resistant allele had great chance to survive upon exposure to pyrethroid only net (Olyset and Interceptor) using cone assays indicating a negative impact of the marker on the efficacy of LLINs as seen for the CYP6P9a/b in *An. funestus* (*5, 11*). Apart from PamaNet 3.0 side, all the other PBO-supplemented nets remained effective including PermaNet 3.0 Top confirming the implication of cytochrome P450s in the observed phenotype (*4, 49, 50*). Considering this negative impact on the efficacy of control tools, it was important to look at the current geographic distribution of the mutation using the diagnostic tool. The study demonstrated high allele frequencies in West africa and central Africa and its potential absence or very low frequency in the South. It should be noted that samples collected from only one country (Malawi) in the South were used in the study in addition to the small sample size. The frequency of this mutation was also found to increase with time (Santchou 2010 vs 2022, Mangoum 2018 vs 2022). This is an indication that the mutation might be under selective pressure (accounted for by the intense use of vector control tools over years) as indicated by a negative Tajima’s D value within the CYP6 cluster. Similar results have been reported with the *kdr* mutations (*16*). The low frequency in *An coluzzii* notably in Ghana may be indicative of recent introgretion event from *An. gambiae* to *An. coluzzi* as previously reported for *kdr* (*51*).

### *CYP6P3*-E205D and *kdr*-L1014F mutations work in concert to reduce bed net efficacy

The *kdr*-L1014F also referred to as the L995F mutation is the main known target site marker involved in pyrethroid and DDT resistance (*9, 25, 26*). The present study is among the first to detect metabolic gene-based markers associated with pyrethroid resistance in *An. gambiae* (*19, 52*) as opposed to *An. funestus* which has had many more (*5, 11, 42, 45, 53*). This study has demonstrated that the *CYP6P3*-E205D and *kdr*-L1014F mutations work together to enhance the ability of mosquitoes to survive exposure to type 1 and 2 pyrethroids using WHO tube and cone assays with pyrethroids implying an additive effect. Our results also showed that Mosquitoes with RR/RS genotype (homozygote resistant for *CYP6P3*-E205D and heterozygote resistant for *kdr* or vice versa) have increased survivorship to pyrethroids exposure than those with the SS/SS genotype. We also showed that resistant field mosquitoes with the *kdr* RR genotype only have lower survivorship compared to those having the *CYP6P3*-205D suggesting that insecticide resistance escalation observed in the field could be multifactorial or polygenic. The same trend was found with pyrethroid only (Olyset and Interceptor) and pyrethroid plus PBO (Permanet 3.0 side) nets using cone assays with double homozygote resistant mosquitoes having better chances to survive than mosquitoes with the susceptible (SS/SS) and other genotypes (Table S6, Fig. 7 **D**). However, mosquitoes succumbed to other PBO nets (Olyset+, Duranet and Permanet 3.0 Top). Mugenzi and colleagues reported similar results in *An funestus* mosquitoes carrying both the 6.5kb structural variant and the CYP6P9a/b mutations (*11*). Our findings indicate that the two markers work in concert to reduce the efficacy of vector control tools specifically pyrethroid only nets and as such constitute a major issue to national control programmes. Another thing that both markers have in common is their predominance or potential origin in west Africa suggesting they may have been selected together by same selective forces.

## Conclusion

This study has successfully deciphered the molecular basis of metabolic resistance to pyrethroids in the major malaria vector *An. gambiae* in West and Central Africa identifying a single amino acid change (E205D) in the *CYP6P3* P450 gene. This allelic variation was shown to confer a greater metabolic ability to the mutant allele to metabolise pyrethroids leading to the design of DNA-based diagnostic assays. This marker is predominant in West and Central Africa and works in concert with the *kdr*-L1014 mutation to reduce the efficacy of pyrethroid-only LLINs. This reliable diagnostic assay will constitute a valuable tool for future vector control programs as it now allows P450-based metabolic resistance to be detected and tracked, assessing cross-resistance to new insecticides, allowing suitable resistance management strategies to be implemented to improve malaria vector control programs in Africa.

## Methods

### Study design

This study was designed to identify genetic variants that confer metabolic-based resistance to pyrethroid insecticides on *An. gambiae* (major malaria vector) and to take advantage of those to develop simple DNA-based diagnostic tools for resistance monitoring in the field. Comparative whole genome sequencing of pyrethroid highly resistant (HR) and highly susceptible (HS) mosquitoes was performed to reveal the regions of interest (signatures of selective sweeps). The highly resistant group was made up of F_0_ field mosquitoes collected from Mangoum (western region of Cameroon) and F_4_ Mangoum_x_Kisumu hybrids that survived 90 min permethrin exposure. The highly susceptible group consisted of the Kisumu laboratory colony and F_4_ Mangoum_x_Kisumu hybrids that succumbed to 3 min permethrin exposure. The genetic crossing was imperative in this study to allow genotype re-segregation as resistance is fixed in Mangoum (*24*). Genome scans were carried out to detect structural variants and point mutations associated with resistance. Because the CYP6 cluster (cluster of P450 cytochromes on CHR 2R) was the main region consistently associated with pyrethroid resistance, genes within this region were Sanger sequenced in HR and HS hybrid mosquitoes to confirm WGS findings. *CYP6P3* was chosen for further investigations because of the 7 associated P450s in the CYP6 locus, it is the main gene to be able to metabolize pyrethroids (*48*)(Muller et al 2008, Edi et al 2014) highlighting the likelihood that polymorphisms in this gene could be most likely drivers of resistance to pyrethroids. To test this hypothesis, *in silico* molecular docking, *in vitro* metabolic assays with recombinant *CYP6P3* proteins, and *CYP6P3* overexpression in transgenic flies were performed to validate the impact of the detected SNP. DNA-based assays (LNA-based probe Taqman and PCR-RFLP assays) were designed to monitor the pyrethroid resistance in mosquitoes collected from across Africa. Finally, we assess the impact of *CYP6P3*-mediated metabolic-based pyrethroid resistance on the effectiveness of long-lasting insecticide-treated bednets using hybrid mosquitoes and standard WHO cone assays.

### Statistical analysis

Statistical analysis was carried out in IBM SPSS statistics v20 with the significance level confidence interval (CI) set at 0.05 and 95% respectively. Fisher’s exact test was used to compare genotype proportions between groups and odds ratio was used to determine the strength of association between the genotype and mortality status for the data obtained from the WHO tube and cone assays. Maps were generated using the quantum geographic information system (QGIS) v3.28.3 and open access shapefiles (https://gadm.org/).

Detailed methodology used for this study is published as supplementary file (see supplementary material and methods).

## SUPPLEMENTARY MATERIALS

Data_file-sheet01 and 2: Polymorphisms and genetic (Excel)

Supplementary material and methods (Pdf)

## Acknowledgements

We are grateful to all those who took part in the collection of archived samples used to access the geographic distribution of the *CYP6P3* marker. Special thanks to Ms Francine Yousseu Sado for drawing the genotypic distribution maps. We are also grateful the following persons; Micareme Tchoupo, Jonathan Chedjoun for their tremendous help with bench work and Nathalie Amvongo for providing the Santchou 2010 DNA samples for genotyping.

## Authors’ contributions

Study conception and design: CSW, JAKO

Sample collection, crossing and bioassays: JAKO, AT, TT, MT

Preparation of Samples for sequencing: JAKO, LM, MK, JH, MW

Poolseq data analysis and interpretation: JAKO, JH, CSW

Functional validation: MK, AM, SSI, MW

Diagnostic tools design and genotyping: JAKO, LM, SN, MW, TT

Paper writing: JAKO, MK, AM, SSI, CSW

Paper proof reading and approval: all authors

## Availability of data and materials

All data generated or analyzed in this study are included in the article and its Additional files. The DNA sequencing data supporting the conclusions of this article are available in the Sequence Read Archive (SRA) of the National Center for Biotechnology Information (NCBI) of the National Institute of Health (NIH), USA, with accession number PRJNA1069339 (http://www.ncbi.nlm.nih.gov/bioproject/1069339). The codes used for data analysis can be made available upon request from the corresponding authors.

## Consent for publication

All authors declare consent for publication.

## Supplementary figures

**Figure S1:**
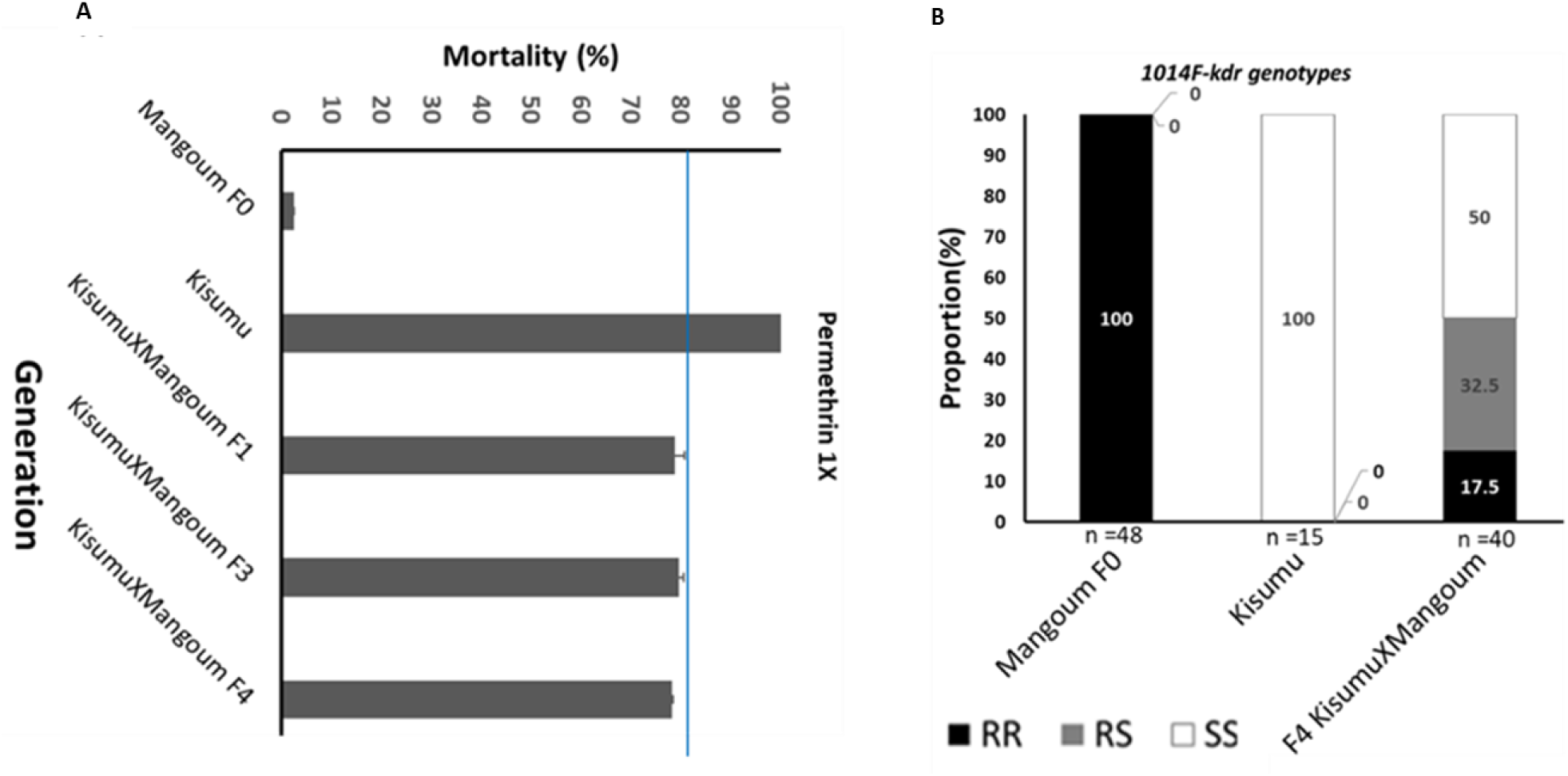
Susceptibility profile and 1014F-kdr genotyping. a) Susceptibility profile of different generations of *An. gambiae* s.s. hybrid (Kisumu_X_Mangoum) mosquitoes to 1X permethrin. Mortality was recorded following one hour of permethrin exposure. Bars represent mean mortality and error bars represent the standard error of the mean (SEM). The bioassay was run with 4 replicates. The blue line corresponds to 80% mortality. b) 1014F-*kdr* genotype proportions in the study hybrid populations.

**Figure S2:**
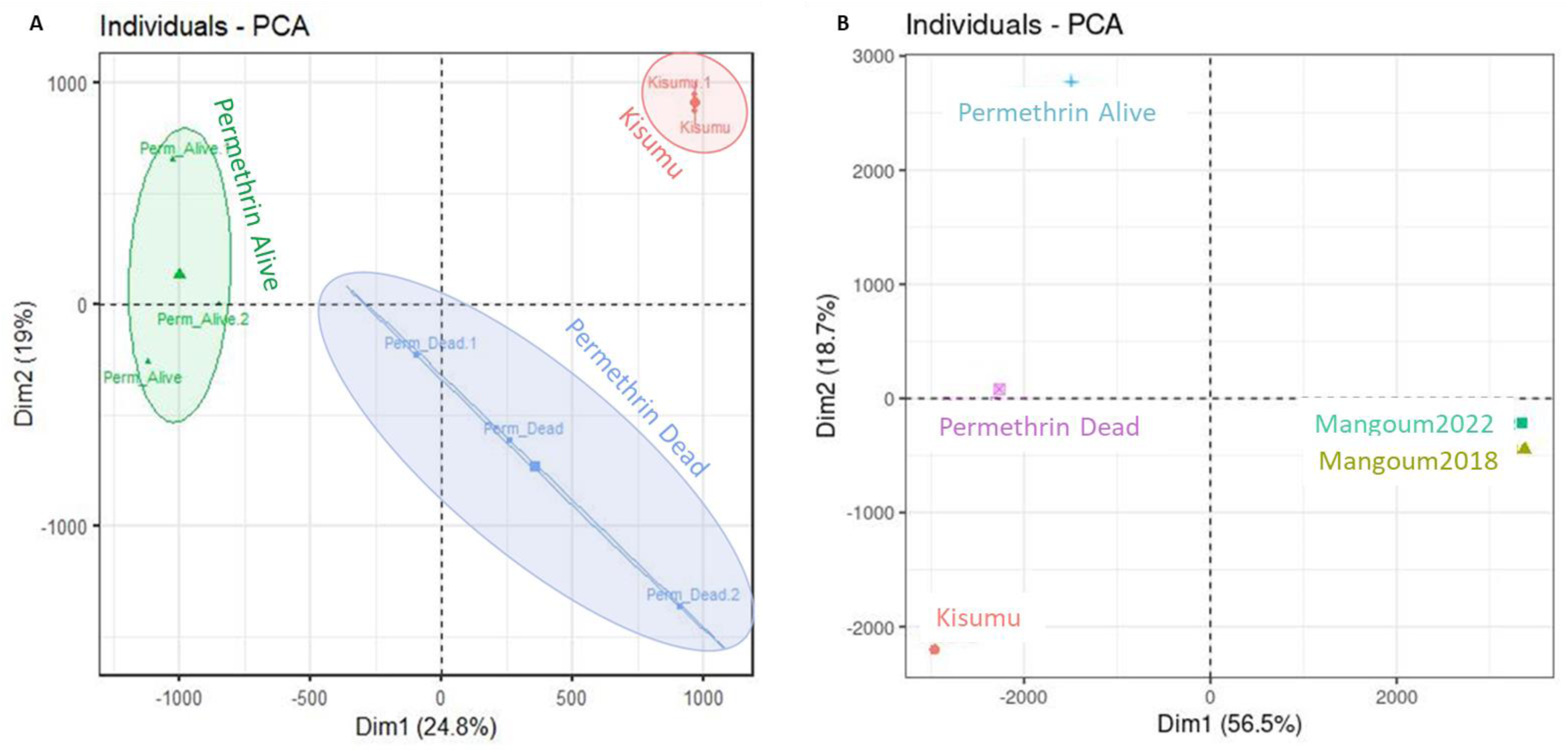
PCA showing clustering of different study populations. Minor allele frequencies (MAF) for Alive, Dead, and Kisumu mosquitoes were used for the PCA using dudi.pca tool of the ade4 package. In (A), the replicates are shown whereas in (B), the replicate for each group are combined and the Mangoum’s pools are also included. Alive and dead mosquitoes were generation from F_4_ Kisumu_X_Mangoum hybrids.

**Figure S3:**
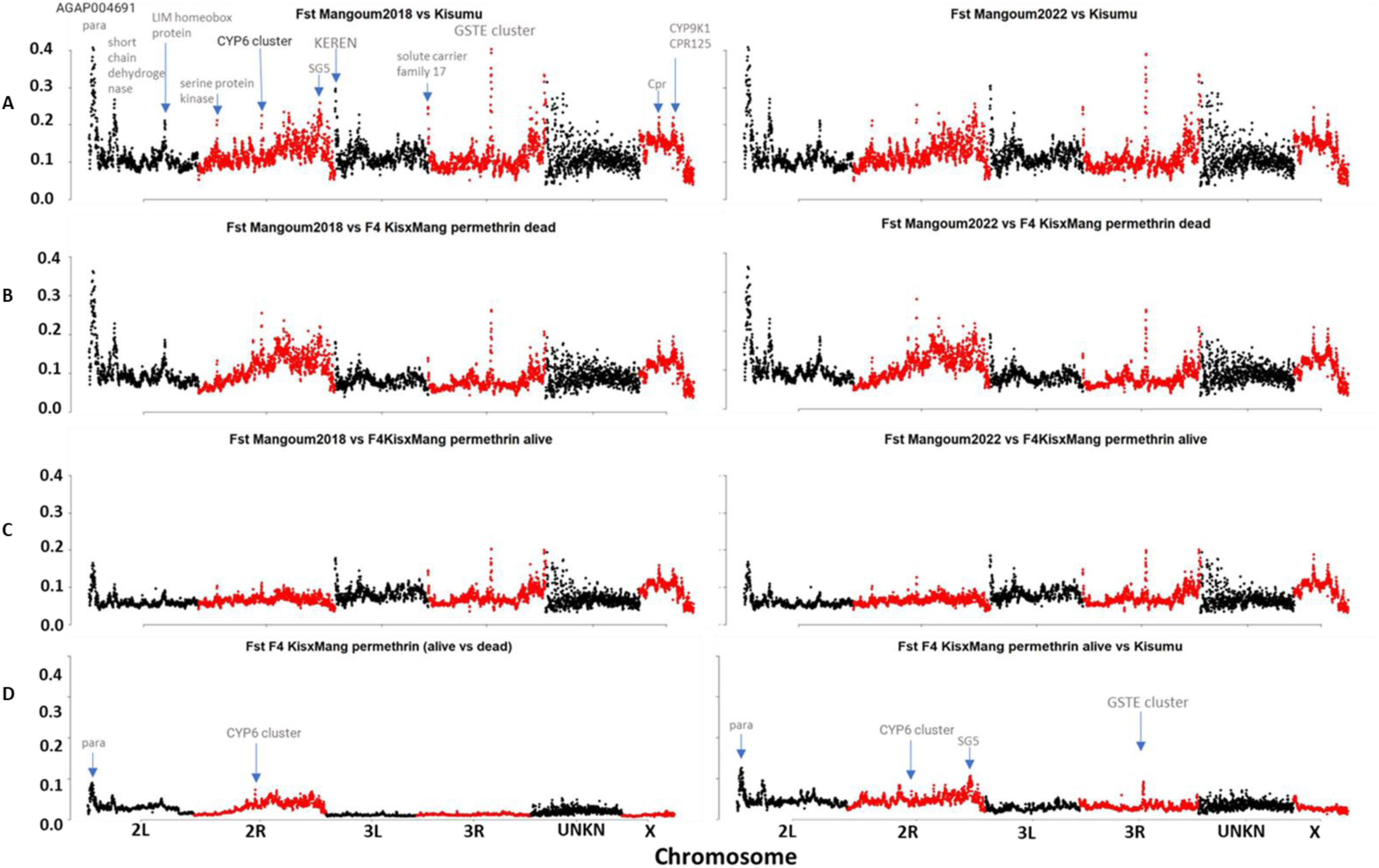
Pair-wise Fst comparison between different study populations. (A) Fst Kisumu versus Mangoum 2018 and Mangoum 2022 respectively. (B) Fst F4 Kisumu_X_Mangoum permethrin dead versus Mangoum 2018 and Mangoum 2022 respectively, (C) Fst F4 Kisumu_X_Mangoum permethrin alive versus Mangoum 2018 and Mangoum 2022 respectively, (D) Fst F4 Kisumu_X_Mangoum permethrin alive versus F4 Kisumu_X_Mangoum permethrin dead and Kisumu respectively. The Fst values were calculated within a window of 100kb and 50kb window steps using winScan and Fst per SNP generated in popoolation2.

**Figure S4:**
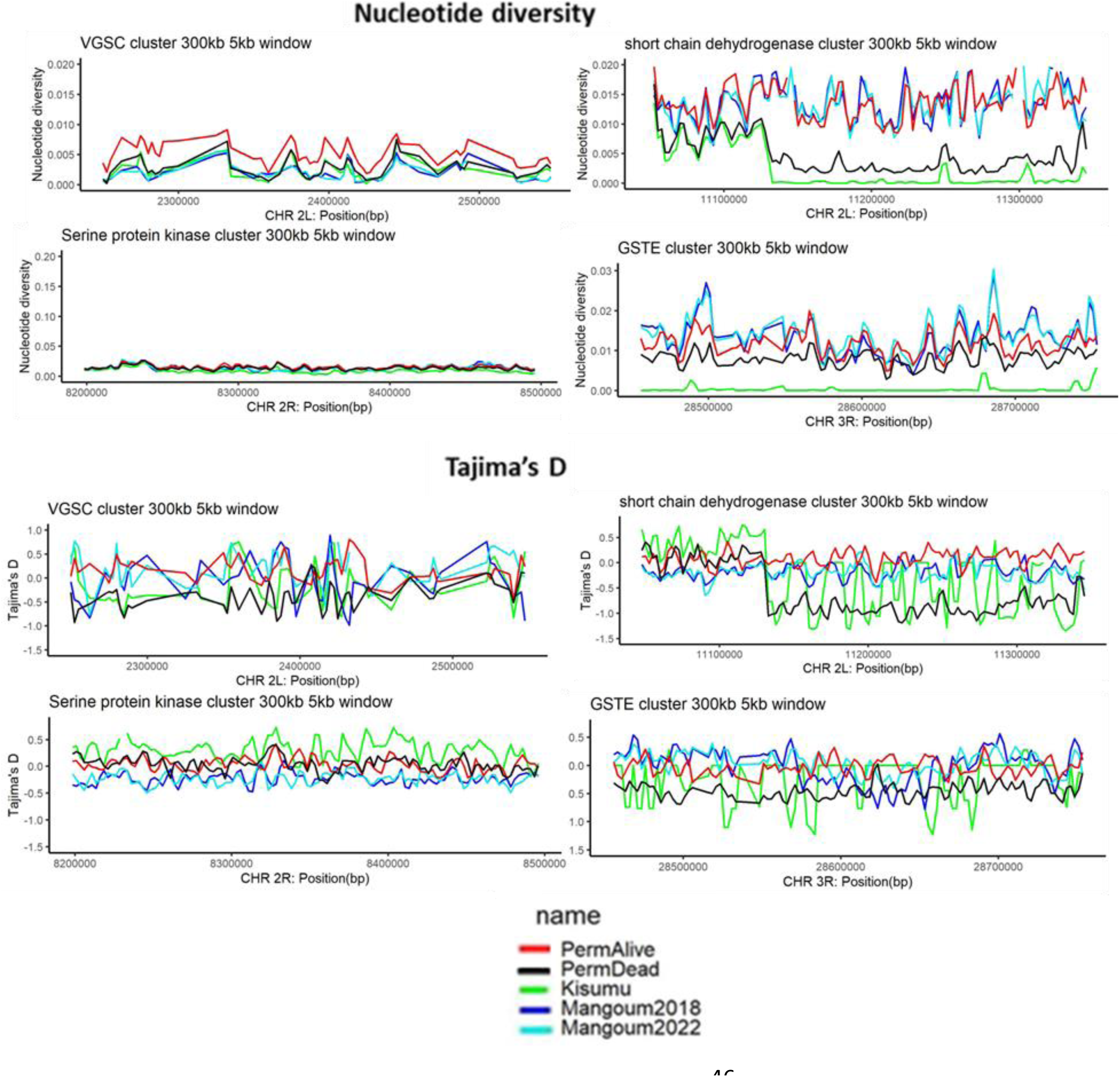
Genetic diversity in regions (clusters 300kb in size) of interest in chromosomes 2L and 2R. Nucleotide diversity (pi) and Tajima’s D was computed using the package popoolation1 within the window of 5000kb and 2500kb window step.

**Figure S5:**
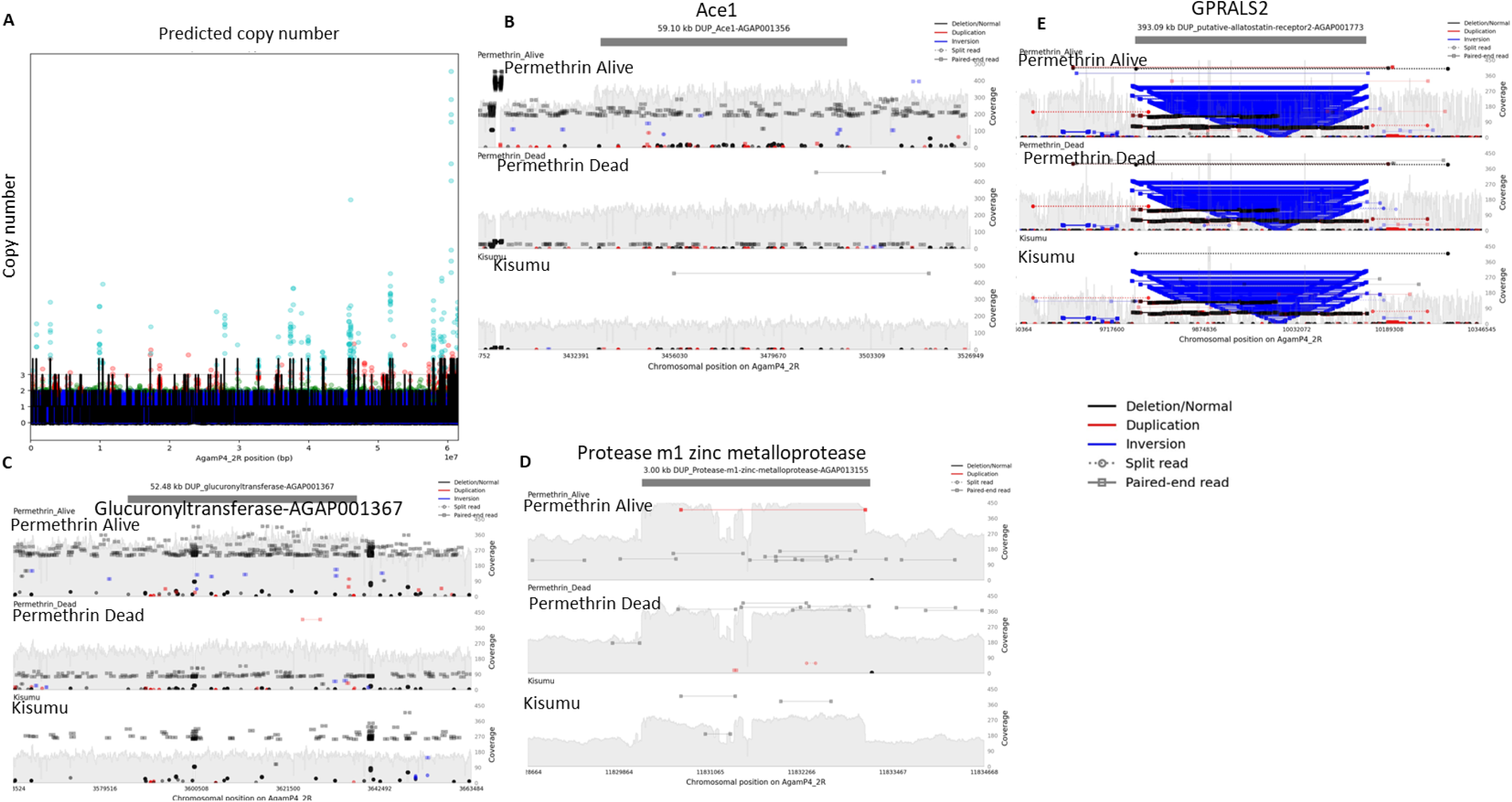
Copy number variation analysis. (A) Predicted copy number across chromosome 2R, coverage visualization (duplications) for (B) Ace1, (c) Glucuronyltransferase-AGAP001367, (D) Protease m1 zinc metalloprotease-AGAP013155 genes between resistant (alive) and susceptible (dead, Kisumu) mosquitoes. (E) Graph showing genomic inversion in putative allostotatin receptor-AGAP001773.

**Figure S6:**
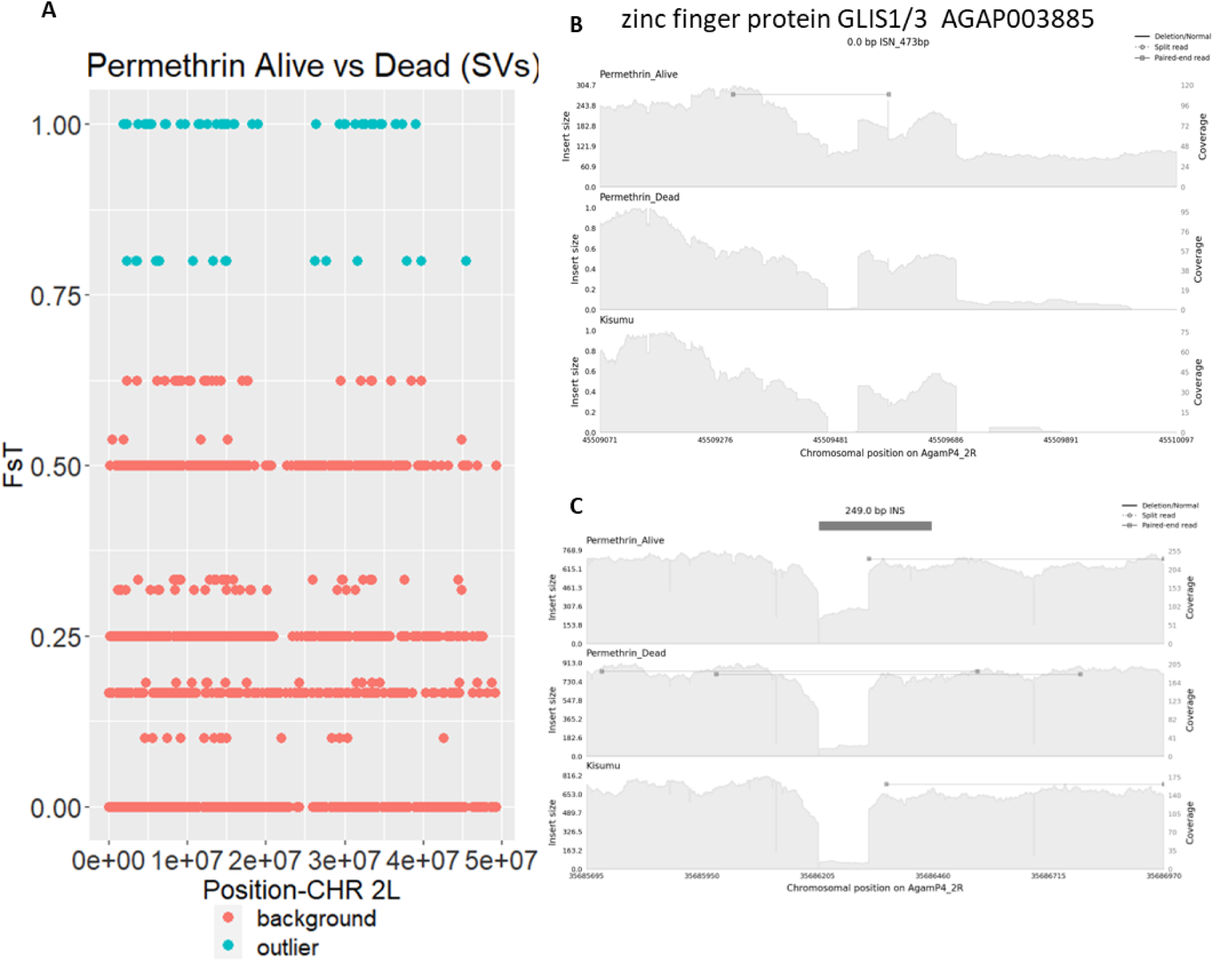
Structural variant analysis. (A) Genetic differentiation between permethrin alive and dead mosquitoes based on structural variants. (B) Samplot graph showing an insertion in the Zinc finger protein (AGAP003885). (C) Samplot graph showing an insertion in a non-coding region on CHR 2R in alive mosquitoes.

**Figure S7:**
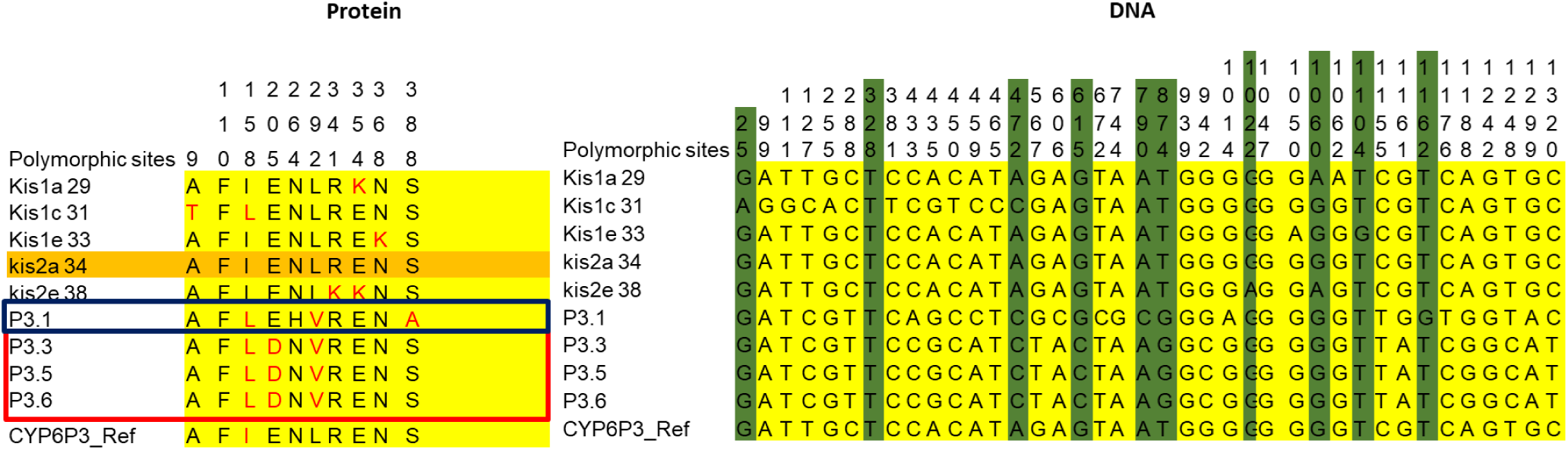
Polymorphism assessment in the CYP6 gene using cDNA from Mangoum and reference Kisumu mosquitoes. cDNAs were cloned in PUAS and Sanger sequenced. Polymorphic sites were detected using MEGA vX. Red and blue boxes indicate the different protein variants. SNP positions in DNA highlighted in green are non-synonymous mutations. Only two haplotypes found in Mangoum clones LHVA and LDV but Kisumu was too diverse.

**Figure S8:**
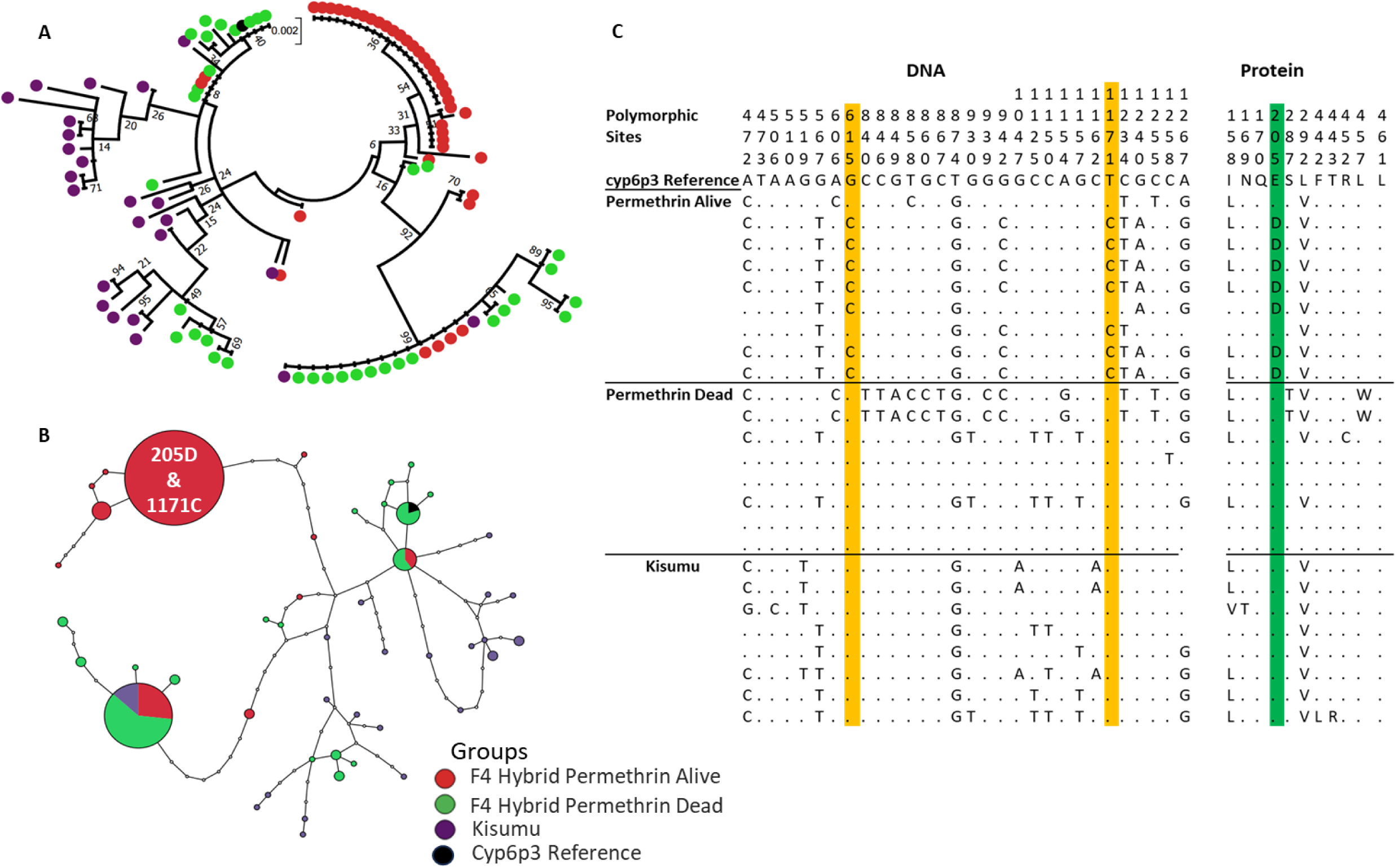
Genetic diversity assessment: (**A**) Maximum likelihood tree of CYP6P3 coding region and intronic regions with F4 Kisumu_X_Mangoum hybrid mosquitoes exposed to permethrin alive (resistant) and dead (susceptible) as well as Kisumu. (**B**) CYP6P3 coding and intronic regions haplotype network (TCS) between Kisumu, susceptible and resistant mosquitoes. (**C**) MEGA-aligned CYP6P3 DNA and protein sequences from resistant, susceptible and Kisumu mosquitoes showing the different polymorphisms. Polymorphic sites highlighted in yellow in genomic sequences represent the G615C and T1171C in the introns with high frequency in resistant mosquitoes. The G615C causes a nonSyn mutation E205D highlighted in green in protein sequences. The tree was built using MEGA v7.0 with 1000 bootstraps.

**Figure S9:**
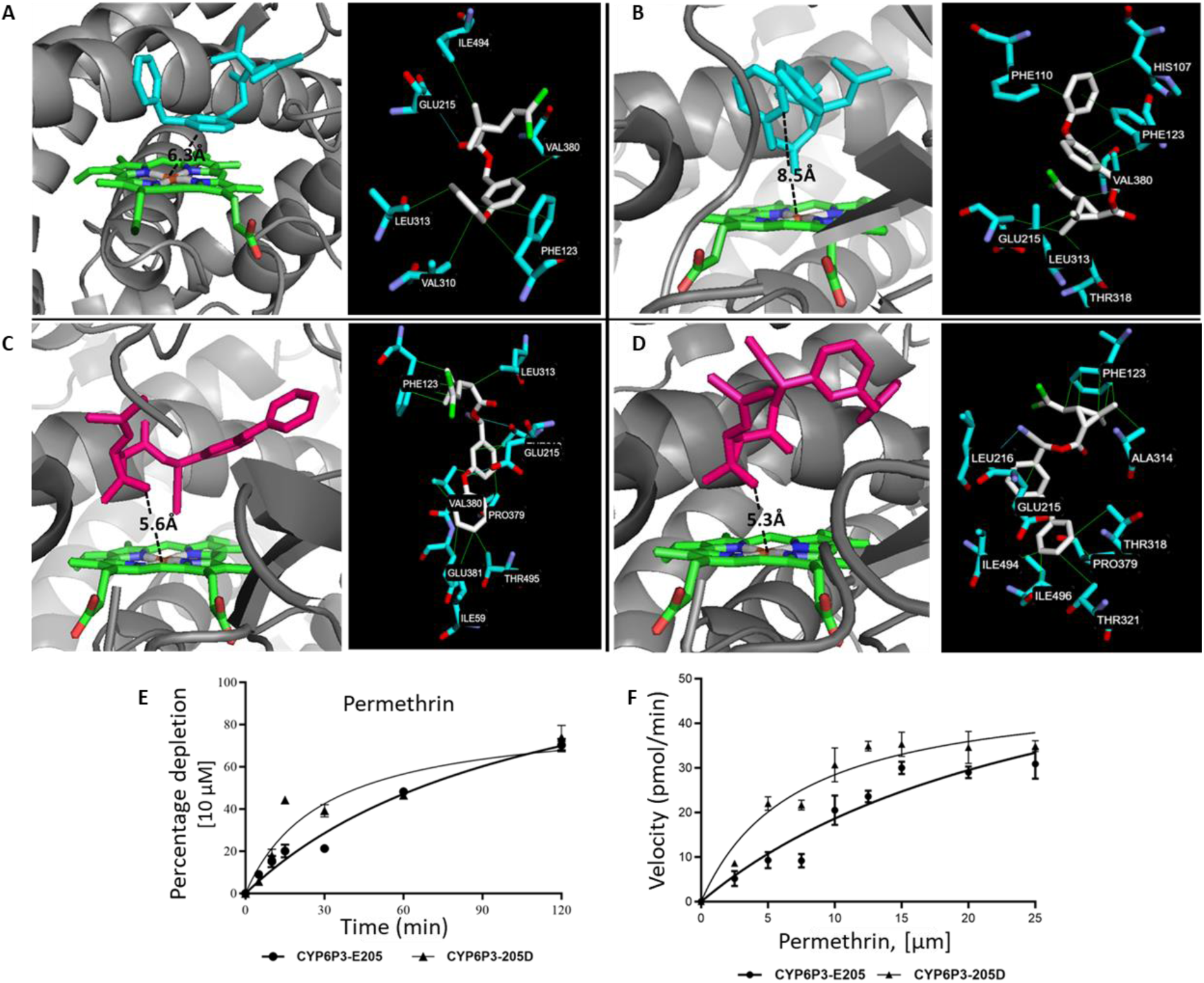
In silico and in vitro functional validation. Predicted binding mode of (A) permethrin in CYP6P3-E205, (B) permethrin in CYP6P3-205D, (C) α-cypermethrin in CYP6P3-E205, and (D) α-cypermethrin in CYP6P3-205D models. CYP6P3 helices are presented in grayscale 50, heme atoms are in stick format and coloured by atoms. Permethrin is in cyan colour and α-cypermethrin in hot pink. Distance between possible sites of metabolism and heme iron are annotated in Angstrom. Insets in black within each panel highlights the predicted interactions between the insecticides and active sites residues contributing to binding energy. (E) Michaelis-Menten plot for the turnover for permethrin the CYP6P3 205D and E205 C alleles. (F) Steady-state kinetic parameters for the metabolism permethrin by CYP6P3 alleles

**Figure S10:**
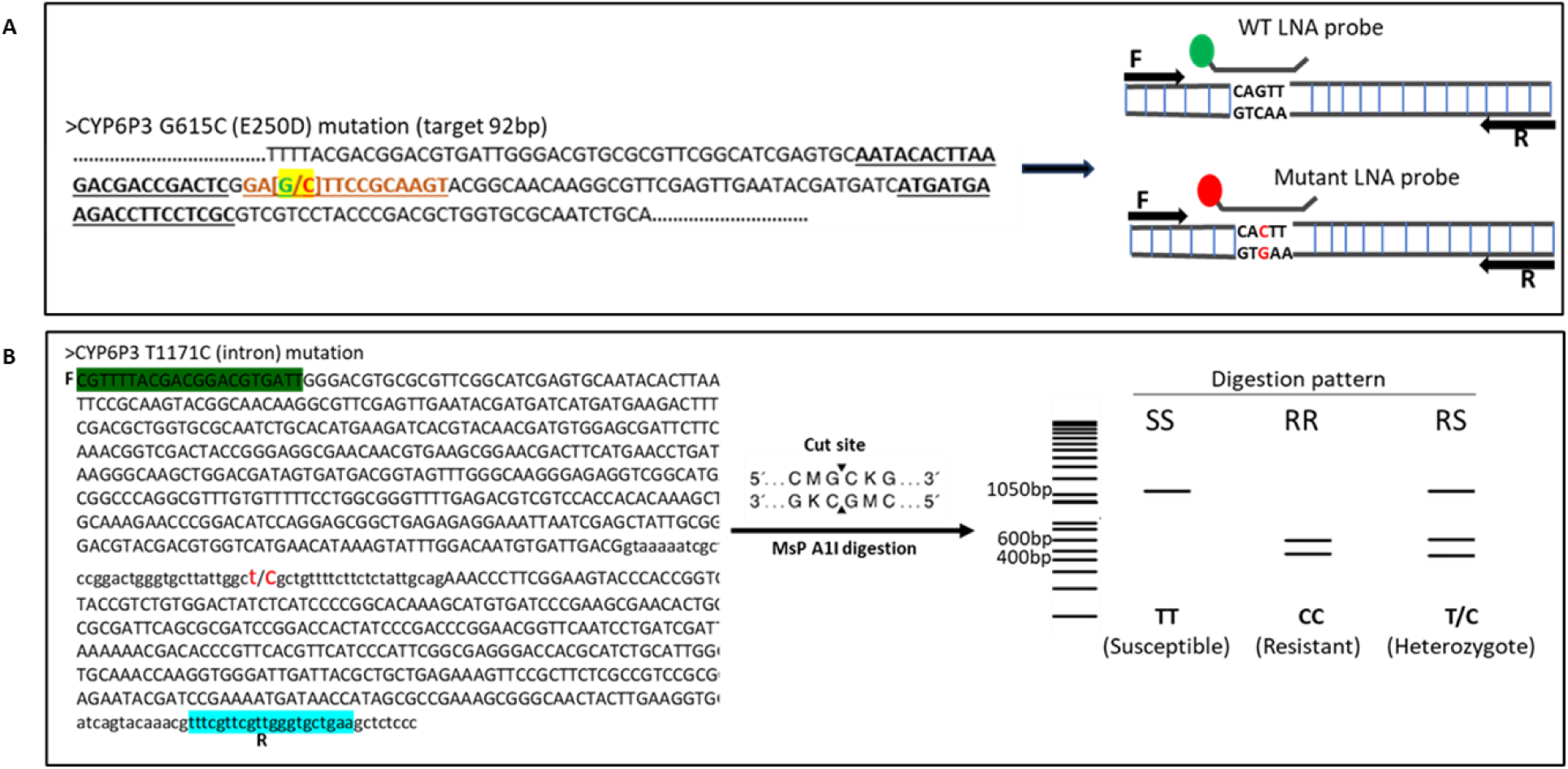
Design of CYP6P3-based diagnostic tools. **(A)** Sequence and LNA Probe Based TaqMan Assay design for the G615C (E250D) Mutation. The probes were designed through IDT and were able to delineate the 615C or 205D (resistant) allele from the susceptible 615G or 205E allele. The mutation is indicated in green and red, with primer binding sites in bold in the sequence and arrows in the LNA tool design (graph with probes indicated). (**B**) Sequence used and MsP Ai digestion pattern prediction for the RFLP assay design using the CYP6P3 T1171C mutation indicated in red colour, the forward and reverse primers are highlighted in green and cyan respectively. The enzyme restriction site and digestion pattern were predicted using a Plasmid editor (APA).

**Figure S11:**
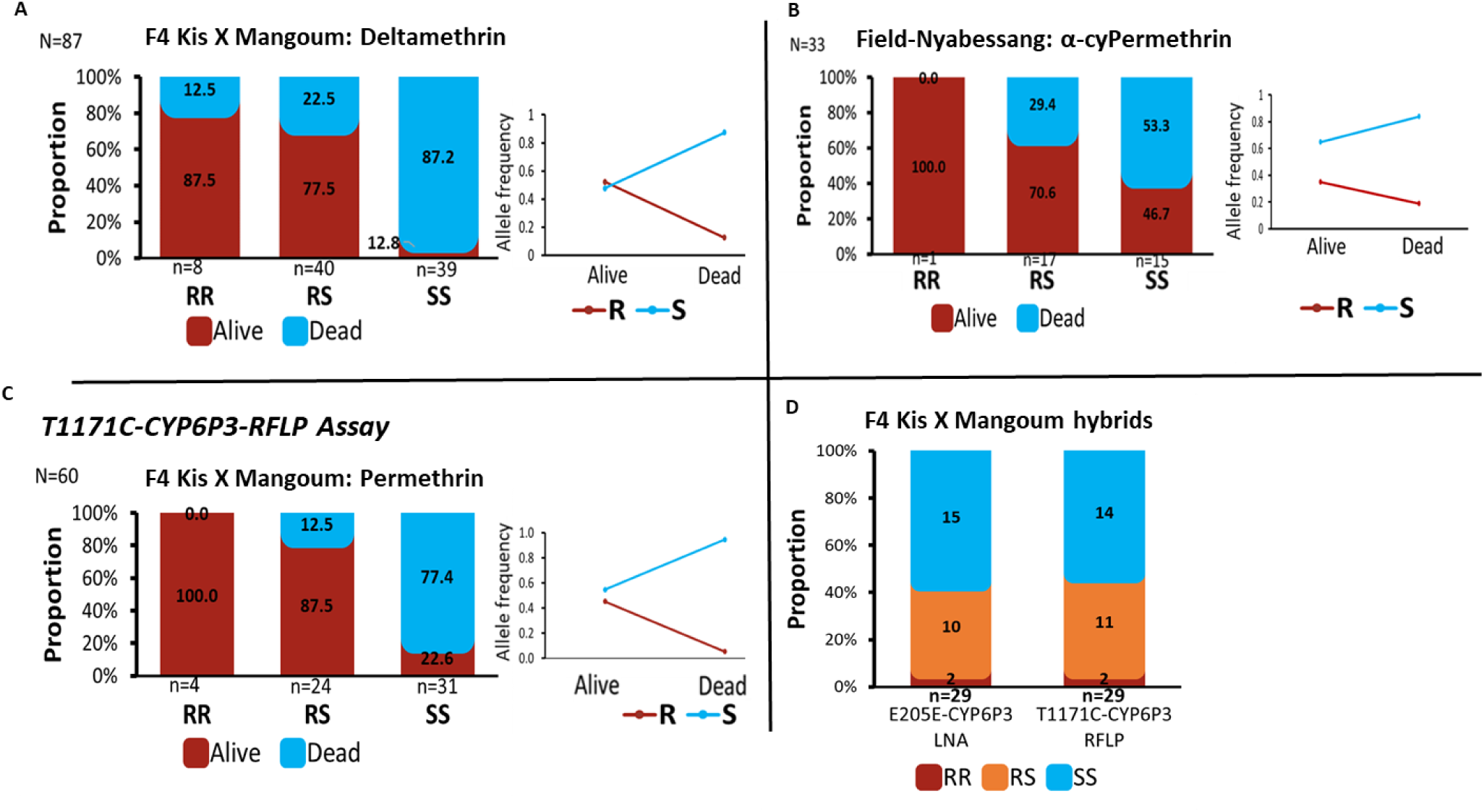
validation of CYP6P3-based diagnostic tools. (A) Association between the CYP6P3E205D markers and deltamethrin resistance using hybrids. (B) Association between the CYP6P3 E205D markers and α-cypermethrin resistance using field mosquitoes. (C) RFLP (T1171C) assay: association between the CYP6P3 markers and permethrin resistance with hybrid mosquitoes using the T1171C-CYP6P3 RFLP assay. (D) Correlation between the E205D-CYP6P3 LNA and T1171C-CYP6P3 RFLP assays.

**Figure S12:**
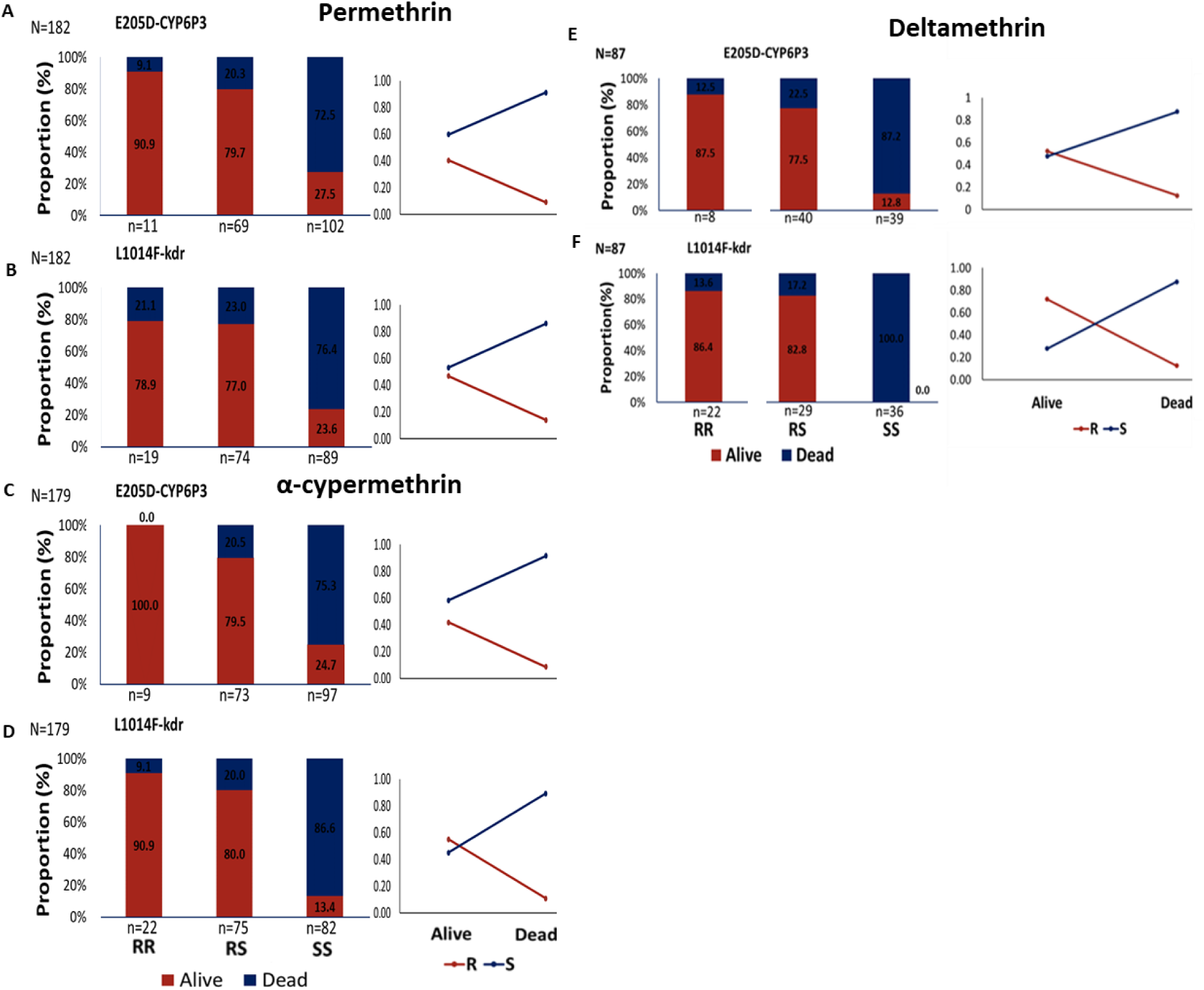
Association between the CYP6P3, kdr markers and pyrethroid resistance with hybrid mosquitoes using the CYP6P3-205D LNA assays. (A) & (B) Assays performed with permethrin-exposed mosquitoes. (C) & (D) Assays performed with α-cypermethrin-exposed mosquitoes. (E) & (F) Assays performed with deltamethrin-exposed mosquitoes. These assays were carried out with DNA extracted from F3/4 Kisumu_X_Mangoum hybrid exposed to different insecticides.

**Figure S13:**
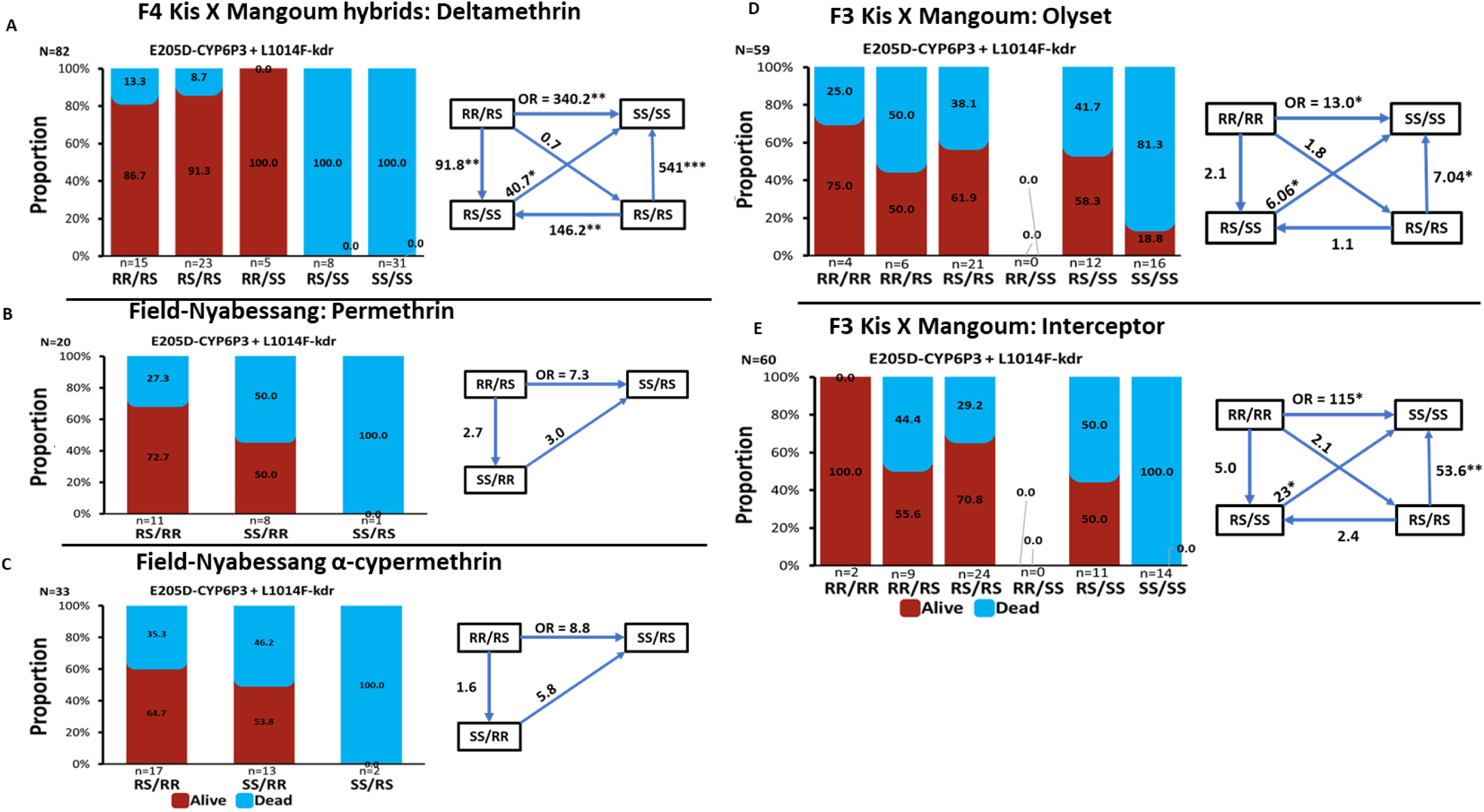
Impact of CYP6P3-E205D and kdr-L1014F markers on pyrethroid insecticide resistance and efficacy of control tools. The figure depicts the frequency distribution of CYP6P3-E205D and kdr-L1014F genotypes between dead and alive *An. gambiae* after exposure to (**A**)1x deltamethrin (F3 Kis X Mangoum hybrids), (**B**)1x permethrin (field samples), (**C**) 1x α-cypermethrin (field samples) using WHO bioassays. (**D**) same as in (a) but with hybrid *An. gambiae* exposed to Olyset (LLIN) using cone assay. (**E**) same as in (d) but with hybrid *An. gambiae* exposed to Interceptor (LLIN) using cone assay. It also gives the odds ratio (OR) and associated significance between different genotypes and the ability to survive exposure to the various insecticides and LLINs. Statistics: “*”=P& 0.05, “**”= P& 0.01, “***”= P &0.001. Kis= Kisumu

**Figure S14:**
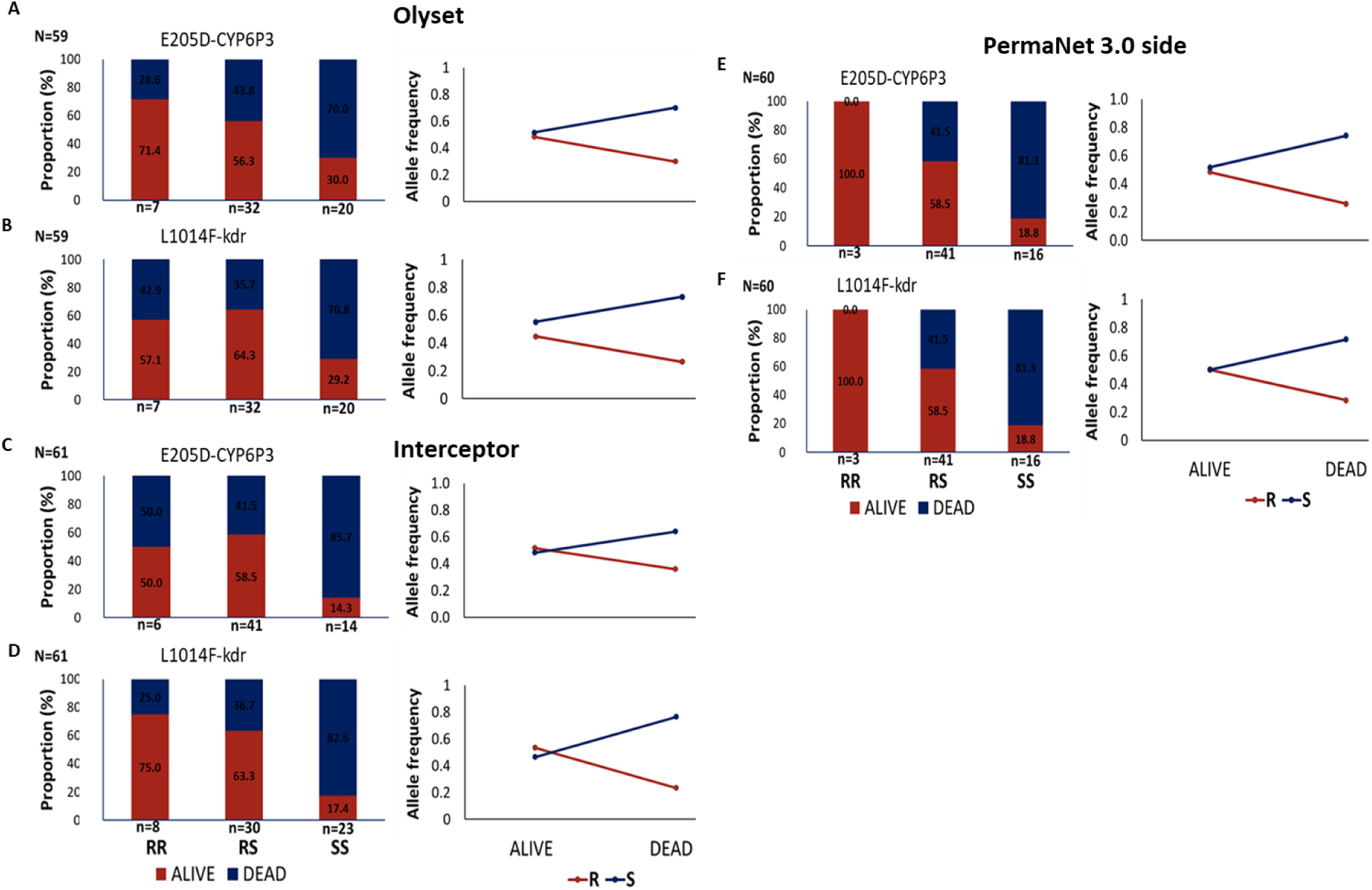
Individual impact of the CYP6P3 and kdr markers on the efficacy of pyrethroid-based bednets using the CYP6P3-205D LNA assays. (A) & (B) Assays performed with Olyset-exposed mosquitoes. (C) & (D) Assays performed with Interceptor-exposed mosquitoes. (E) & (F) Assays performed with PermaNet 3.0 side-exposed mosquitoes. These assays were carried out with DNA extracted from F3/4 Kisumu_X_Mangoum hybrid exposed to different bednets.

**Figure S15:**
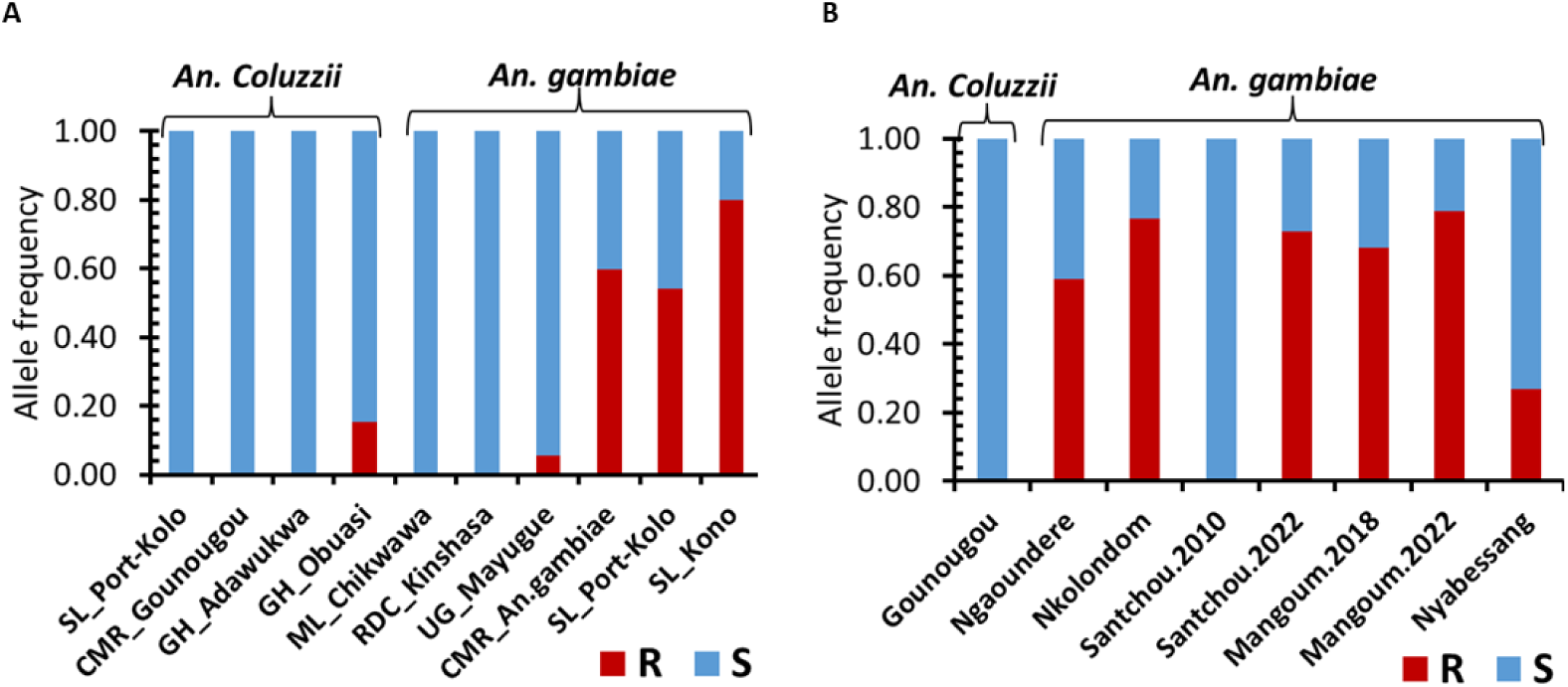
Geographic distribution of the CYP6P3 E205D mutation in Africa following genotyping of archived samples. (**A**) Histogram showing the frequency of the CYP6P3 E205D mutation across Africa. (**B**) Histogram showing the frequency of the CYP6P3 E205D mutation across Cameroon. SL=Sierra Leone, CMR=Cameroon, GH=Ghana, ML=Malawi, RDC=Republic Democratic of Congo, An=Anopheles. Samples used were collected between 2021 and 2022 (see Figure 8).

## Supplementary tables

**Table S1:**
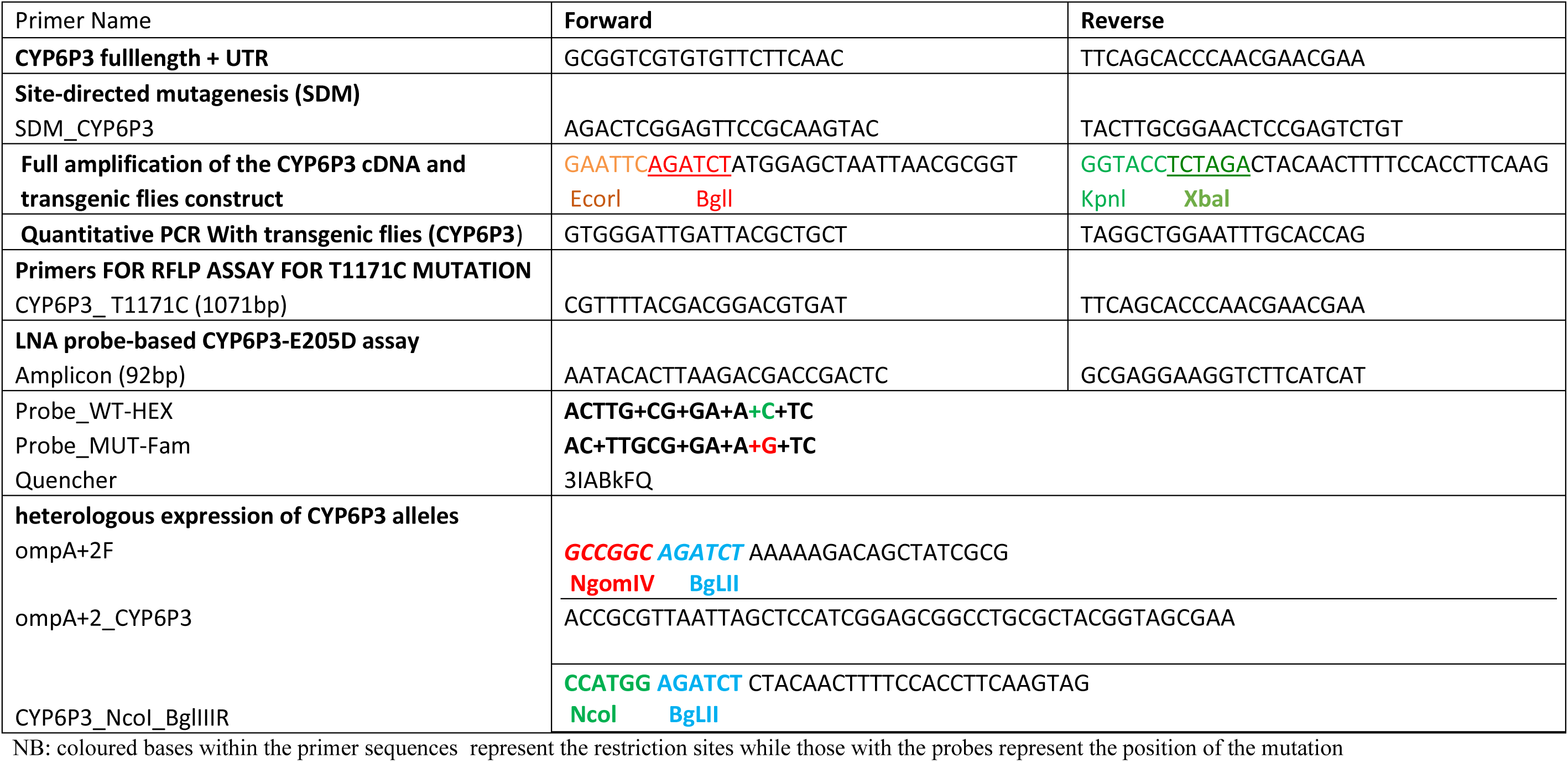
List of primers and enzymes used in the study

**Table S2:**
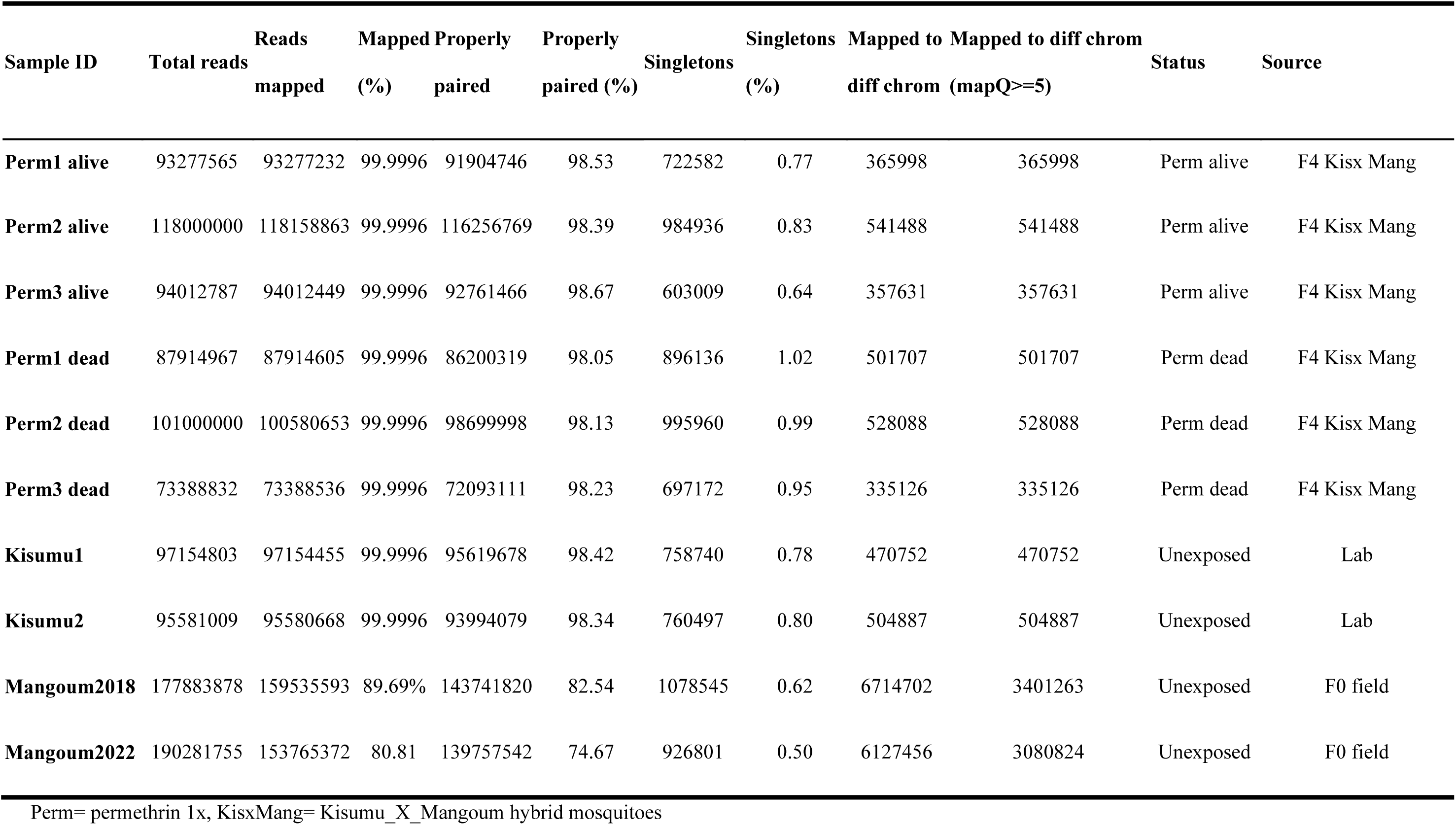
Bam files statistics for all samples after mapping to the reference genome

**Table S3:** CYP6P3 Genetic diversity parameters

| Gene | Samples | # of sequences (2n) | Length | # of polymorphic sites | NonSyn | Syn | Pi ($\pi$) | # of Haplotypes | Haplotype diversity | Tajima's D | Fu & Li's F |
| --- | --- | --- | --- | --- | --- | --- | --- | --- | --- | --- | --- |
|  | **cDNA** |  |  |  |  |  |  |  |  |  |  |
|  | Mangoum F<sub>0</sub> | 4 | 1530 | 23 | 5 | 18 | 0.0049 | 2 | 0.5 | -0.84729 | -0.8668 |
|  | Kisumu | 12 | 1530 | 25 | 7 | 18 | 0.00343 | 10 | 0.955 | -1.63394 | -2.0986 |
| **Genomic coding and intronic regions** |  |  |  |  |  |  |  |  |  |  |  |
|  | F<sub>4</sub> hybrid alive | 40 | 1057 | 21 | na | na | 0.00529 | 11 | 0.686 | 0.44311 | 1.49505 |
|  | F<sub>4</sub> hybrid dead | 36 | 1057 | 32 | na | na | 0.01054 | 18 | 0.921 | 1.40383 | 1.965* |
|  | Kisumu | 24 | 1057 | 39 | na | na | 0.01087 | 21 | 0.989 | 0.08155 | 1.3165 |
| **Genomic coding region** |  |  |  |  |  |  |  |  |  |  |  |
|  | F<sub>4</sub> hybrid alive | 40 | 978 | 19 | 3 | 16 | 0.00514 | 11 | 0.686 | 0.41126 | 1.45769 |
|  | F<sub>4</sub> hybrid dead | 36 | 978 | 28 | 5 | 23 | 0.01 | 18 | 0.921 | 1.38145 | 1.9299* |
|  | Kisumu | 24 | 978 | 34 | 9 | 25 | 0.00966 | 21 | 0.989 | -0.17836 | 1.19257 |
| **Genomic intronic region** |  |  |  |  |  |  |  |  |  |  |  |
|  | F<sub>4</sub> hybrid alive | 40 | 79 | 2 | na | na | 0.00721 | 3 | 0.666 | 0.3977 | 0.76794 |
|  | F<sub>4</sub> hybrid dead | 36 | 79 | 4 | na | na | 0.01728 | 3 | 0.641 | 1.01693 | 1.20033 |
|  | Kisumu | 24 | 79 | 5 | na | na | 0.02582 | 8 | 0.841 | 1.51701 | 1.46751 |

**Table S4:**
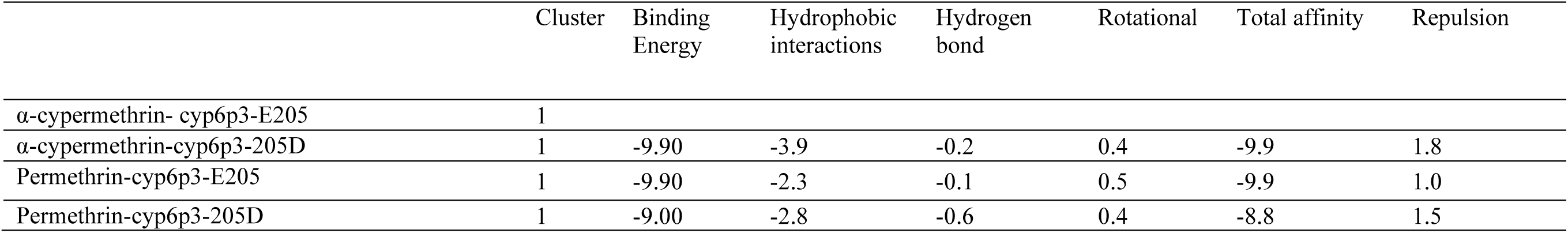
Predicted binding parameters of the top-ranked docking mode of various insecticides in CYP6P3-E205 and CYP6P3-205D models The binding energy and the energetic contribution to binding energy (repulsion, hydrophobic interactions, hydrogen bonds, total affinity) are all in kcal/mol.

|  | Cluster | Binding Energy | Hydrophobic interactions | Hydrogen bond | Rotational | Total affinity | Repulsion |
| --- | --- | --- | --- | --- | --- | --- | --- |
| α-cypermethrin- cyp6p3-E205 | 1 | | | | | | |
| α-cypermethrin-cyp6p3-205D | 1 | -9.90 | -3.9 | -0.2 | 0.4 | -9.9 | 1.8 |
| Permethrin-cyp6p3-E205 | 1 | -9.90 | -2.3 | -0.1 | 0.5 | -9.9 | 1.0 |
| Permethrin-cyp6p3-205D | 1 | -9.00 | -2.8 | -0.6 | 0.4 | -8.8 | 1.5 |

**Table S5:** Correlation between the CYP6P3-205D/kdr-L1014F genotypes and ability to survive pyrethroid insecticide exposure using WHO tube assay.

| **E205D-CYP6P3** | **Genotypes**<br>**CYP6P3/kdr** | **Odds ratio (OR)** | **95%Confidence**<br>**interval (CI)** | **P value** |
| --- | --- | --- | --- | --- |
| Permethrin | RR vs SS | 26.4 | 3.2-216.07 | P = 0.0023 |
|  | RR vs RS | 2.5 | 0.3-21.5 | P = 0.3917 |
|  | RS vs SS | 10.4 | 5.0-21.5 | P < 0.0001 |
|  | R vs S | 6.7 | 3.05-14.9 | P < 0.0001 |
| Alphacypermethrin | RR vs SS | 57.0 | 3.2-1015.8 | P = 0.0059 |
|  | RR vs RS | 5.0 | 0.28-91.3 | P = 0.2744 |
|  | RS vs SS | 11.8 | 5.6-24.4 | P < 0.0001 |
|  | R vs S | 7.3 | 3.3-16.1 | P < 0.0001 |
| Deltamethrin | RR vs SS | 43.4 | 4.3-432.2 | P = 0.0013 |
|  | RR vs RS | 2.0 | 0.2-18.7 | P = 0.5318 |
|  | RS vs ss | 21.3 | 6.4-70.9 | P < 0.0001 |
|  | R vs S | 7.9 | 3.8-16.1 | P < 0.0001 |
| **L1014F-kdr** |  |  |  |  |
| Permethrin | RR vs SS | 12.1 | 3.6-40.5 | P = 0.0001 |
|  | RR vs RS | 1.1 | 0.3-3.8 | P = 0.8583 |
|  | RS vs SS | 10.9 | 5.2-22.5 | P < 0.0001 |
|  | R vs S | 5.4 | 2.7-10.8 | P < 0.0001 |
| Alphacypermethrin | RR vs SS | 64.5 | 13.2-315.3 | P < 0.0001 |
|  | RR vs RS | **2.5** | 0.5-11.8 | P = 0.2496 |
|  | RS vs SS | 25.8 | 11.02-60.4 | P < 0.0001 |
|  | R vs S | 9.9 | 4.7-20.7 | P < 0.0001 |
| Deltamethrin | RR vs SS | 406.7 | 19.9-8282.6 | P = 0.0001 |
|  | RR vs RS | 1.3 | 0.3-6.2 | P = 0.7264 |
|  | RS vs ss | 325.1 | 17.2-6151.2 | P = 0.0001 |
|  | R vs S | 19.3 | 9.1-40.8 | P < 0.0001 |
| **E205D-CYP6P3 + L1014F-kdr** |  |  |  |  |
| Permethrin | RR/RS vs RS/RS | 1.25 | 0.2-7.1 | P = 0.8016 |
|  | RR/RS vs RR/SS | 1.4 | 0.11-18.6 | P = 0.7909 |
|  | RR/RS vs RS/SS | 5.9 | 1.2-28.9 | P = 0.0263 |
|  | RR/RS vs SS/SS | 64.8 | 12.5-334.1 | P < 0.0001 |
|  | RS/RS vs RR/SS | 1.1 | 0.1-11.5 | P = 0.9156 |
|  | RS/RS vs RS/SS | 4.7 | 1.5-14.5 | P = 0.0056 |
|  | RS/RS vs SS/SS | 51.8 | 15.7-171.1 | P < 0.0001 |
|  | RR/SS vs RS/SS | 4.2 | 0.47-37.9 | P = 0.1988 |
|  | RR/SS vs SS/SS | 45.7 | 4.8-430.4 | P = 0.0008 |
|  | RS/SS vs SS/SS | 10.8 | 4.2-27.8 | P < 0.0001 |
| Alphacypermethrin | RR/RS vs RS/RS | 2.6923 | 0.3- 24.6 | P = 0.3801 |
|  | RR/RS vs RR/SS | 3.5 | 0.2-64.6 | P = 0.3999 |
|  | RR/RS vs RS/SS | 21 | 2.5-172.2 | P = 0.0046 |
|  | RR/RS vs SS/SS | 260.4 | 28.7-2358.1 | P < 0.0001 |
|  | RS/RS vs RR/SS | 1.3 | 0.13-13.1 | P = 0.8240 |
|  | RS/RS vs RS/SS | 7.8 | 2.5-24.1 | P = 0.0004 |
|  | RS/RS vs SS/SS | 96.72 | 26.3-355.8 | P < 0.0001 |
|  | RR/SS vs RS/SS | 6 | 0.6-54.7 | P = 0.1121 |
|  | RR/SS vs SS/SS | 74.4 | 7.4-745.6 | P = 0.0002 |
|  | RS/SS vs SS/SS | 12.4 | 4.1-37.6 | P < 0.0001 |
| Deltamethrin | RR/RS vs RS/RS | 0.6 | 0.07-4.9 | P = 0.6511 |
|  | RR/RS vs RR/SS | 0.5 | 0.02-11.9 | P = 0.6624 |
|  | RR/RS vs RS/SS | 91.8 | 3.9-2153.7 | P = 0.0050 |
|  | RR/RS vs SS/SS | 340.2 | 15.3-7571.6 | P = 0.0002 |
|  | RS/RS vs RR/SS | 0.7 | 0.03-18.7 | P = 0.8793 |
|  | RS/RS vs RS/SS | 146.2 | 6.3-3372.8 | P = 0.0019 |
|  | RS/RS vs SS/SS | 541.8 | 24.7-11854.1 | P = 0.0001 |
|  | RR/SS vs RS/SS | 187 | 3.2-10885.6 | P = 0.0116 |
|  | RR/SS vs SS/SS | 693 | 12.4-38727.9 | P = 0.0014 |
|  | RS/SS vs SS/SS | 40.7647 | 2.1-812.7 | P = 0.0152 |

**Table S6:** Correlation between the E205D-CYP6P3/L1014F genotypes and ability to survive LLINs using WHO cone assay.

|  | Genotypes | Odds ratio (OR) | 95%Confidence interval (CI) | P value |
| --- | --- | --- | --- | --- |
| **E205D-CYP6P3** |  |  |  |  |
| Olyset | RR vs RS | 1.9 | 0.3-11.5 | P = 0.4646 |
|  | RR vs SS | 5.4 | 0.8-36.3 | P = 0.0820 |
|  | RS vs SS | 2.7 | 0.8-9.2 | P = 0.0924 |
|  | R vs S | 2.05 | 1.15-3.6 | P = 0.0145 |
| Interceptor | RR vs RS | 0.7 | 0.12-3.9 | P = 0.6938 |
|  | RR vs SS | 6 | 0.6-53.6 | P = 0.1090 |
|  | RS vs SS | 8.5 | 1.6-42.8 | P = 0.0098 |
|  | R vs S | 1.98 | 1.1-3.5 | P = 0.0185 |
| PermaNet 3 side | RR vs RS | 4.9 | 0.5-46.04 | P = 0.1607 |
|  | RR vs SS | 13.7 | 1.38-136.2 | P = 0.0254 |
|  | RS vs SS | 2.7 | 0.89-8.6 | P = 0.0780 |
|  | R vs S | 2.6 | 1.45-4.7 | P = 0.0015 |
| **L1014F-kdr** |  |  |  |  |
| Olyset | RR vs RS | 0.7 | 0.1-3.9 | P = 0.7270 |
|  | RR vs SS | 3.2 | 0.5-18.3 | P = 0.1848 |
|  | RS vs SS | 4.3 | 1.3-14.1 | P = 0.0136 |
|  | R vs S | 2.2 | 1.2-3.9 | P = 0.0085 |
| Interceptor | RR vs RS | 1.7 | 0.3-10.1 | P = 0.5397 |
|  | RR vs SS | 14.2 | 2.06-98.1 | P = 0.0070 |
|  | RS vs SS | 8.2 | 2.2-30.3 | P = 0.0016 |
|  | R vs S | 3.7 | 2.05-6.9 | P < 0.0001 |
| PermaNet 3 side | RR vs RS | 5 | 0.2-103.08 | P = 0.2972 |
|  | RR vs SS | 27 | 1.1-654.4 | P = 0.0427 |
|  | RS vs SS | 6.1 | 1.5-24.8 | = 0.0113 |
|  | R vs S | 2.5 | 1.4-4.6 | P = 0.0016 |
| **E205D-CYP6P3 /L1014F-kdr** |  |  |  |  |
| Olyset | RR/RR vs RR/RS | 3.0 | 0.2-47.9 | P = 0.4373 |
|  | RR/RR vs RS/RS | 1.8 | 0.16-20.94 | P = 0.6207 |
|  | RR/RR vs RS/SS | 2.1 | 0.17-27.1 | P = 0.5561 |
|  | RR/RR vs SS/SS | 13.0 | 0.9-172.9 | P = 0.0521 |
|  | RR/RS vs RS/RS | 0.6 | 0.09-3.8 | P = 0.6024 |
|  | RR/RS vs RS/SS | 0.7 | 0.09-5.1 | P = 0.7377 |
|  | RR/RS vs SS/SS | 4.3 | 0.5-33.1 | P = 0.1577 |
|  | RS/RS vs RS/SS | 1.16 | 0.27-4.9 | P = 0.8400 |
|  | RS/RS vs SS/SS | 7.04 | 1.5-32.6 | P = 0.0126 |
|  | RS/SS vs SS/SS | 6.1 | 1.1-33.2 | P = 0.0378 |
| Interceptor | RR/RR vs RR/RS | 4.1 | 0.15-108.9 | P = 0.4002 |
|  | RR/RR vs RS/RS | 2.1 | 0.1-50.2 | P = 0.6358 |
|  | RR/RR vs RS/SS | 5.0 | 0.2-122.7 | P = 0.3243 |
|  | RR/RR vs SS/SS | 115.0 | 1.8-7307.7 | P = 0.0251 |
|  | RR/RS vs RS/RS | 0.5 | 0.1-2.5 | P = 0.4107 |
|  | RR/RS vs RS/SS | 1.3 | 0.2-6.7 | P = 0.7947 |
|  | RR/RS vs SS/SS | 28.1 | 1.2-619.9 | P = 0.0345 |
|  | RS/RS vs RS/SS | 2.4 | 0.6-9.5 | P = 0.2037 |
|  | RS/RS vs SS/SS | 53.6 | 2.7-1033.4 | P = 0.0083 |
|  | RS/SS vs SS/SS | 23.0 | 1.1-465.2 | P = 0.0410 |
| PermaNet 3 side | RR/RR vs RR/RS | 1.6 | 0.04-58.23 | P = 0.7782 |
|  | RR/RR vs RS/RS | 3.3 | 0.14-77.9 | P = 0.4457 |
|  | RR/RR vs RS/SS | 5.5 | 0.2-130.3 | P = 0.2891 |
|  | RR/RR vs SS/SS | 95 | 1.4-6088.1 | P = 0.0319 |
|  | RR/RS vs RS/RS | 2.6 | 0.25-27.4 | P = 0.4099 |
|  | RR/RS vs RS/SS | 4.4 | 0.4-47.5 | P = 0.2172 |
|  | RR/RS vs SS/SS | 57 | 1.9-1693.5 | P = 0.0195 |
|  | RS/RS vs RS/SS | 1.6 | 0.5-5.56 | P = 0.4059 |
|  | RS/RS vs SS/SS | 28.1 | 1.4-535.7 | P = 0.0267 |
|  | RS/SS vs SS/SS | 17.2 | 0.8-337.1 | P = 0.0610 |

## Suppl material and methods

### Material and Methods

#### Sample collection

Mosquito larvae were collected from different breeding sites using the dipping method, at Mangoum, an agricultural setting in the Western region of Cameroon between April 2021 and January 2022. These F0 samples are referred to in this study as Mangoum2022. Also, blood fed adult mosquitoes, resting indoor were aspirated indoors early in the morning (6:00 AM-10:00 AM) from the walls and the roof of houses across the village using electric aspirators.

Both larvae and blood fed female mosquitoes were transported to the insectary for of the Centre for Research in Infectious Diseases (CRID) in Yaoundé for morphological identification (1, 2) and rearing. A recent characterization of this population has shown that the An. gambiae ss was predominant and exhibited high level of resistance to pyrethroids (10X) while also resistant to DDT, carbamates and organophosphates (3).

#### Mosquito rearing and crossing

The adult F0 resistant field mosquitoes and susceptible Kisumu lab colony were forced to lay eggs (Morgan et al., 2010), and the eggs allowed to hatch in water. Larvae were fed with Tetramin™ baby fish food till pupation, and the pupae transferred individually in falcon tubes. Upon emergence, female and male adults from Kisumu and field mosquitoes were sorted out and transferred in a common cage for mating and generation of the F1 hybrid Kisumu_x_Mangoum lines. The hybrid line was then propagated till the 4th filial generation (F4) that was used for the tests outlined below.

#### WHO tube and cone bioassays

For susceptibility testing and selection of highly susceptible and highly resistant mosquitoes, F3 and F4 female hybrid Kisumu_x_Mangoum lines were subjected to a modified WHO protocol for adults (4) using different insecticide classes. The following insecticides were tested: pyrethroid type 1 (0.75% permethrin), pyrethroid type 2 (0.05% α-cypermethrin and 0.05% deltamethrin), and a carbamate (0.1% bendiocarb). Prior to selection, a couple of assays were carried out with different exposure times to determine the LT20 (exposure time resulting in 20% mortality) and LT80 (exposure time resulting in 80% mortality) used for the bioassays. Briefly, 3 to 4-d old female hybrid female mosquitoes (n = 20-25) were transferred into tubes containing insecticide-embedded papers and exposed for a length of time equal to either the LT20 and TL80. Mosquitoes that died at LT20 were considered highly susceptible (Dead), while those that survived at LT80 were defined as highly resistant (Alive). The Dead (LT20) and alive (LT80) mosquitoes were stored individually at -20°C in 80% ethanol for subsequent DNA extraction and molecular analyses.

The F3 female hybrid mosquitoes were also exposed to different bed nets to assess their bio-efficacy using cone assays following the WHO’s instructions (4). The following nets were tested: pyrethroid only nets; Olyset (2% permethrin), Interceptor (200 mg/m2 α-cypermethrin), Duranet (270 mg/m2 α-cypermethrin), PermaNet 3.0 side (85 mg/m2 ±25% deltamethrin), Olyset Plus (2% permethrin + 1% PBO), and PermaNet 3.0 top panel (85 mg/m2 ±25% deltamethrin + PBO).

Ten female mosquitoes (3-4 d old) were aspirated into plastic cones sealed with the impregnated net and exposed for 3 min. They were immediately transferred to paper cups and fed with 10% sucrose solution and knock down was monitored after 1 h and mortality after 24 h. Untreated net was used as control. Dead and alive mosquitoes were kept in silica gel at 4°C for molecular work.

#### DNA extraction and quantification

DNA extraction was carried out using Qiagen DNeasy kit (Qiagen, Hilden, Germany) following the manufacturers’ instructions for samples intended for NGS while the Livak protocol (5) was used for samples intended for genotyping. DNA quantification was performed using Nanodrop (Thermo Scientific, MA, USA) and the fluorimetric Qubit™ dsDNA BR Assay Kits (Thermo Scientific, MA, USA) in Qubit® 2.0 Flurometer (Life Technologies, CA, USA). DNA was extracted from the lab Kisumu colony, F0 An gambiae non-exposed samples (Mangoum 2022), archived mosquitoes collected in 2018 (Mangoum, 2018), and insecticide-exposed Kisumu_x_Mangoum hybrid female mosquitoes (alive and dead individuals).

#### Whole genome Pooled DNA sequencing (PoolSeq)

Experimental Procedures: Prior to sequencing, DNA was extracted from individual mosquitoes. pools of 50 samples each were generated and RNAse A (QIAGEN, UK) treated to get rid of RNA contaminants according the manufacturers’instructions. Pooling was done at a concentration of 400ng/sample (NanoDrop) to maximize yield following RNAse A treatment. RNAse treated DNA pools were sent to Novogene (Cambridge, UK) for library preparation and sequencing (see suppl material). Quality control for size distribution was done using an Agilent Bioanalyzer. Libraries were sequenced on an Illumina Novaseq platform resulting in the production of paired-end reads in the fastq format.

Sequence analysis: Upon receiving the fastq files, reads’ quality was assessed using FastQC v0.11.5. Low quality reads were trimmed with Trimmomatic V0.32 using the default parameters. Cleaned reads were mapped to the reference Anopheles gambiae genome version 59 (https://vectorbase.org/common/downloads/release-59/AgambiaePEST/fasta/data/VectorBase-59_AgambiaePEST_Genome.fasta) using BWA mem (Version: 0.7.17). Duplicates were marked and removed from the mapped reads using the Picard tools. The coverage was computed using the coverage option of the samtools package (version 1.13). Popoolation2 v. 1.201 was used to compute genetic differentiation per SNP between populations. The FST was then computed per window of 100kb and 50kb step-size using WindowScanr (version 0.1). Using Popoolation2-generated minor allele frequencies, the principal component analysis was done using ade4 and factoextra packages. Popoolation (v.1.2.2) was used to assess the genetic diversity in the regions of interest (regions surrounding the major peaks). The indels were filtered and ignored using the identify-genomic-indel-regions.pl and filter-pileup-by-gtf.pl scripts in Popoolation. As population genetic estimators are sensitive to sequencing errors (highly covered regions), the data was subsampled to a uniform coverage of 15 using the subsample-pileup.pl script before nucleotide diversity and Tajima’s D calculation. This was done within a window-size of 10kb and a step-size of 5kb. These parameters were computed within a region of 300kb covering each of the main peaks obtained with popoolation2 (Fst). To identify non-synonymous mutations likely associated with the phenotype, VarScan.v2.3.9 was used for SNP calling with the default parameters. Then, annotation was performed with SnpEffect. The genome-wide Fst values were visualized using the package qqman v.0.1.8, Pi and tajima’s D with ggplot2. All these packages were implemented in the R environment. Permethrin-exposed hybrid mosquitoes and field collected Mangoum 2018 and 2022 samples were used for Poolseq analyses.

#### Assessment of genetic diversity of full length CYP6P3

Following Poolseq analyses, further attention was given to the CYP6P cluster on 2R chromosome, notably the CYP6P3 gene (AGAP002865). Primers were designed to amplify a 2500bp fragment covering the CYP6P3 gene in permethrin-exposed F4 Kisumu_X_Mangoum hybrid dead and alive mosquitoes (Table S1). The amplification was done using Kappa Taq DNA amplification kit as recently described (3). The following cycling parameters were used: initial denaturation 95°C for 5min, 35 cycles of denaturation 95°C for 30s, annealing 69°C for 45s, extension 72°C for 1 min, 30s, and a final extension 72°C for 10min. The PCR products were then treated with ExoSAP-IT PCR Product Cleanup Reagent (ThermoFisher Scientific, MA, USA) and sent for Sanger sequencing. The DNA sequences were aligned using Bioedit for polymorphisms detection. Genetic diversity assessment was done using DnaSP V6.12 (6). The package TCS was used for haplotype network analysis. For phylogenetic tree construction, fasta sequences were aligned using mafft-7.029 with the auto option, MEGAX (7) was used for phylogenetic trees inference by maximum likelihood with 1000 bootstraps.

#### Homology Modelling and Docking Simulations

To investigate differences in the abilities of the CYP6P3 alleles to interact with pyrethroid insecticides, 3D models of the P450 alleles were created using Easy Modeller (8), using CYP3A4 structure (PDB: 1TQN, ∼32.5% identity for the two sequences) (9) as a template. A total of 20 models were generated for each sequence and assessed externally using Errat version 2.0 (10), to identify the best model from statistical patterns of non-bonded interactions between different atom types. Overall quality scores were 57.82% and 62.45% for the highest ranked models, respectively for the CYP6P3-E205 and CYP6P3-205D models. Virtual α-cypermethrin structure (PubChemID: 93357) was downloaded from PubChem (https://pubchem.ncbi.nlm.nih.gov/compound/101618973) and 1R-cis permethrin (ZINC01850374) was retrieved from the library in ZINC12 database (11). Docking simulations were carried out using Blind Docking Server, available at http://bio-hpc.ucam.edu/webBD/index.php/entry (12). For each ligand, 30 binding poses were generated and sorted according to binding energy and conformation in the model’s active site. Figures were prepared using the PyMOL 1.7 (13). Non-bonded interactions were predicted using protein-ligand interaction profiler (14).

### Assessing the effect of the E205D mutation on the activity of CYP6P3 using heterologous expression and metabolism assays

#### Amplification and cloning of CYP6P3-205D Mangoum allele

The full-length amino acids coding sequence of CYP6P3 harbouring the novel 205D mutation was copied from cDNA prepared from the wild Mangoum population of An. gambiae. PCR amplification was carried out using Phusion Hot Start II Taq polymerase (Thermo Fisher Scientific, MA USA), following previously published protocols (15–17). The full length sequence was amplified fused to the amino terminal of the bacterial ompA leader sequence with downstream ala+pro linker according to the method of (18). Products amplified using Primers in Table S1 and which were cloned into PjET1.2 vector were digested using the NgomIV and BgLII restriction enzymes. Purified fragments were ligated into the linearised pCWori+ vector digested with the same restriction enzymes. The construct was co-transformed in E. coli JM109 competent cells (Thermo Fisher Scientific, MA USA), together with plasmid carrying the An. gambiae cytochrome P450 reductase (CPR) pACYC_Ag_CPR (17, 19).

#### Site-directed mutagenesis of Mangoum CYP6P3-205D into CYP6P3-E205

A previously established primer extension PCR protocol (20, 21) was used to introduce a single mutation E205 into Mangoum allele (harbouring D205 residue). The Mangoum construct pB13::ompA+2-CYP6P3-205D in Pjet1.2 vector was used as the template for amplification using Phusion HotStart II Taq Polymerase (Thermo Fisher Scientific, MA, USA). The PCR mix comprised 0.12 μl Hot start II polymerase, 0.2 μl dNTP (25 mM), 3 μl of HF Buffer, 1 μl of the plasmid template, 0.51 μl each of the 10 mM of the forward and reverse mutagenesis primers (Table S1). The PCR reaction was carried out in a final Volume of 15 μl. Thermocycling conditions comprised initial denaturation for 10 s at 98 °C, followed by 35 cycles with annealing at 62 °C and an extension of 5 min at 72 °C. A final extension at 72 °C for 20 min. The PCR product (3 μl) was run on 1% agarose gel to confirm amplification and the rest of the amplicon digested using DpnI restriction enzyme (New England, Biolabs, USA). The digested amplicon was purified using the Qiaquick kits (Qiagen, Hilden Germany) and the product transformed in DH5α cells (Thermo Fisher Scientific, USA). Positive colonies were miniprepped and sequenced to confirm the mutation. Plasmid with the mutation was subsequently used in digestion with NgomIV and BgLII restriction enzymes and the fragments ligated into pCWori+ linearized with the same enzyme to create a pCW-ori+-CYP6P3-E205 construct. This construct was co-transformed together with the plasmid bearing An. gambiae CPR, as explained above, and subsequently used for expression.

#### Heterologous expression of the recombinant CYP6P3 alleles

The expression and purification of recombinant CYP6P3 was conducted according to the previous protocols (17, 22). Expression started with a starter culture in which 2ml LB supplemented with ampicillin and chloramphenicol was inoculated with positive colonies containing the plasmids. A 200ml terrific broth medium (Alpha Teknova, Hollister, USA) was supplemented with the starter cultures and allowed to shake at 200 rpm and 37 °C until the cultures were ready for induction with an absorbance of ∼0.8-1.0. Cultures were induced with 1 mM isopropyl β-D-1-thiogalactopyranoside (IPTG) and 0.5 mM δ-aminolevulinic acid and maintained until expression was obtained.

#### In vitro metabolism assays with insecticides and the determination of steady state kinetic parameters

In vitro metabolism assays were conducted using methods previously described (15–17). Assays were conducted with pyrethroids (permethrin, deltamethrin and α-cypermethrin). The membranes co-expressing recombinant P450s CYP6P3-E205 and CYP6P3-205D proteins together with AgCPR at 0.1 μM concentration were constituted in 0.1M potassium phosphate buffer at pH 7.4, with cytochrome b5 protein, at the ratio 1:10. Reactions were started by adding the mixture containing the recombinant proteins, buffer and 10 μM insecticides with NADPH regeneration buffer. The reaction mixes were incubated at 30 °C for 2 h with shaking at 1200 rpm. Reactions were quenched with the addition of 200 μl ice-cold acetonitrile and the mix spun at 15, 000 rpm for 20 min. One hundred microliters of the supernatants were loaded into vials for injection in the HPLC for analysis. For the HPLC running conditions, pyrethroids were analysed with mobile phase 70% acetonitrile and 30% water, flow rate of 1 mL/min, and detection wavelength of 226 nm for 30 min. All the insecticides were detected on the Hypersil GOLD™ C18 Selectivity HPLC Column 250 x 4.6mm, particle size 5um (Thermo Fisher Scientific).

To determine the reaction rates and the steady state kinetic parameters of permethrin and α-cypermethrin for the two alleles, reactions were carried out initially for the turnover by keeping the enzyme and substrates concentrations constant (50 picomole of recombinant P450s and 10 μΜ insecticides) but varying the time of the reaction starting from 5 min to 120 min. For the steady state kinetic parameters, reactions conducted using 50 pmol of the recombinant P450s and the reaction rates were monitored for 15 min with varying substrate concentrations between 2.5-25μM as described earlier (16). Using the plots of substrates concentrations against the velocities of the reactions and fitting that into the Michaelis-Menten module, the steady state kinetic parameters of Km and Vmax were determined for each allele.

### Assessing the impact of CYP6P3 E205D mutation on insecticide resistance using transgenic **D. melanogaster flies**

#### Creation of transgenic flies expressing the variant allele of CYP6P3

To confirm whether the overexpression of CYP6P3-205D can confer cross resistance to pyrethroid and other insecticide classes, transgenic flies overexpressing the CYP6P3 were generated using the GAL4-UAS system following the protocols previously described (23, 24). The CYP6P3-205D allele was amplified using Phusion Taq polymerase with the allele specific primers (Table S1) and ligated to the D. melanogaster expression vector pUASattB pre-digested with BglII and KpnI restriction enzymes (New England Biolabs, Hitchin, UK). Ligation was achieved using Thermofisher Ligation kit (Fermentas, Burlington, Ontario, Canada) following the manufacturer’s recommendations. The generated construct was used to transform DH5α cells using and plasmids midiprepped using HiSpeed Plasmid Midi Kit (QIAGEN, Manchester, United Kingdom) creating pUASattB::CYP6P3 constructs. The pUASattB::CYP6P3 constructs was sent to the fly genetic department of Cambridge University (Genetic Services, MA, USA) for injection into flies.

Constructs were transformed into the germ line of PhiC31 D. melanogaster strain carrying the attP450 docking site on 25C6 chromosome 2 [y w M(eGFP, vas-int, dmRFP)ZH-2A; P{CaryP}attP40]. The transgenic flies containing 205D-CYP6P3 were successful generated with red eyes and no carly wing phenotypes and delivered at the Centre for Research in Infectious Disease (CRID) for further analysis. They were delivered alongside a control line (D. melanogaster which received an empty pUASattb vector), with white eyes and no carly wing as phenotype (72). Microinjection and balancing of UAS stock to remove integrase was carried out by Fly Facility (Cambridge, UK) generating a UAS-CYP6P3 transgenic line. Ubiquitous expression of the transgene in adult F1 progeny (experimental group) was obtained after crossing virgin females from the GAL4-Actin driver strain Act5C-GAL4, BL25374 [y[1] w[*]; P{Act5C-GAL4-w}E1/CyO, 1;2] (Bloomington, IN, USA) with the UAS-CYP6P3 males. Similarly, adult F1 control progeny (control group) with the same genetic background as the experimental group but without the CYP6P3 insert were obtained by crossing virgin females from the driver strain Act5C-GAL4 and UAS recipient line males with white eyes (not carrying the pUASattB-CYP6P3 insertion). The GAL4 line was purchased from Bloomington Stock Centre (http://flystocks.bio.indiana.edu/). The flies were reared at 25°C in plastic vials with flies’ food. To confirm overexpression of CYP6P3 in the experimental group three replicates of F1 females were used for qRT-PCR, using a previously established protocol (25). Total RNA was extracted, and cDNA was synthesized and the relative expression levels of the transgene were measured using the experimental F1 progeny as well as the controls. The primers used are provided in Table S1.

#### Insecticide contact assays

Three to five days old F1 adult female flies expressing the CYP6P3-205D allele and control flies were exposed for 24h to different insecticide classes including pyrethroids (permethrin (4%); deltamethrin (2%), α-cypermethrin (0.007%), organochloride (DDT 4%), Carbamates (propoxur (0.1%), bendiocarb (0.007) using impregnated papers, prepared in acetone and Dow Corning 556 Silicone Fluid (BHD/Merck, Hesse, Germany). These papers were rolled and introduced into 45 cc plastic vials to cover the entire wall. The vials were plugged with cotton soaked in 10% sucrose. For each insecticide, 4 pools of 20-25 of experimental and control lines were exposed and the mortality was recorded after 1h, 2h, 3h, 6h, 12h and 24h exposure.

### Design of DNA-based CYP6P3 diagnostic tests

Upon assessing the genetic diversity of the CYP6P3 gene, two SNPs (G615C->E205D and T1171C within the introns) found to be in the same haplotype and associated with pyrethroids resistance after SANGER sequencing were exploited for the development of an LNA-based and RFLP-based diagnostic tools, respectively. For the G615C->E205D SNP, a primer set flanking a short 92bp region covering the mutation and two locked nucleic acid (LNA) based probes conjugated to FAM and HEX for the mutant and wild type allele, respectively were commercially designed and acquired from IDT (Table S1). PCR amplification was carried out in a final volume of 10µL containing 1 µmole of each probe, 2 µmoles of each primer in 1× PrimeTime Master Mix (IDT) or 1× Luna Universal qPCR Master Mix (NEB) and 1 µL of genomic DNA. Amplification was achieved using the 2 fast step settings with a hold at 95°C for 10 min and 40 cycles of 95°C for 10s denaturation and 60°C for 45s extension.

The T1171C RFLP assay was developed by determining the presence of restriction sites in the SNP region using a plasmid editor v2.0.30 (APE) software. Primer sets spanning that region was designed (Table S1). For this assay, DNA amplification was done using the Ben Taq kit (Beneficial Bio, Cameroon) with a final forward and reverse primers concentration of 0.34µM, 1x ben Taq mix and 1uL of genomic DNA. A total of 35 amplification cycles were achieved using T100 thermal cycler (BIO-RAD) with an initial denaturation at 95°C for 5min, denaturation at 95°C for 30s, annealing 62.5°C for 30s and extension at 72°C for 45s. After amplification, PCR products were then digested with the restriction enzyme MsPAI (New England Biolabs) at a final concentration of 0.12 U/µL in 1x MsPAI buffer for 4h at 37°C following the manufacturer’s instructions. The digests were resolved on 1.5% agarose (SIGMA, St. Louis, USA).

### Assessment of correlation between the CYP6P3-E205D and pyrethroid resistance

To assess the ability of the CYP6P3-E205D assay to detect mosquitoes resistant to both type I and II pyrethroids, F4 Kisumu_X_Mangoum hybrid female mosquitoes were exposed to 0.75% permethrin, 0.05% deltamethrin and 0.05% α-cypermethrin using WHO bioassays. Alive and dead mosquitoes were genotyped and comparison made to estimate correlation between CYP6P3 genotypes and resistance phenotypes. Moreover, field mosquitoes from Nyabessang in Cameroon, in 2021 were also tested and genotyped, to confirm if this marker confers resistance to permethrin and deltamethrin in the field populations.

### Assessment of the impact of the CYP6P3-E205Dallele on the efficacy of bed nets

To assess the impact of the CYP6P3-E205Dallele on the performance of insecticide-treated nets, the CYP6P3-E205Dmarker was also genotyped in F3 Kisumu_X_Mangoum hybrid An. gambiae mosquitoes exposed to different insecticide-treated nets, including PermaNet 3.0, Olyset, Olyset Plus, Interceptor, DuraNet using cone assays (26). Briefly, 5 replicates of 10 F3 females (2–5 d old) were placed in plastic cones attached to 30cm x 30cm pieces of nets, and an insecticide-free net as a control. After 3 min of exposure, the mosquitoes were immediately transferred to holding paper cups and provided with cotton soaked in a 10% sugar solution. Mortality was recorded 24 h post-exposure. The bed nets were tested Alive and dead mosquitoes were genotyped and comparison made to establish the impact of CYP6P3 genotypes and bed net efficacy.

### Africa-wide geographic field distribution of An. gambiae CYP6P3-205Dmarker

To determine the geographic distribution of the CYP6P3-E205D in field mosquitoes, archived F0 An gambiae samples collected from different African countries (Sierra Leone, Ghana, Cameroon, Democratic Republic of Congo (DRC), Uganda and Malawi) were genotyped using the CYP6P3-E205DLNA probe-based tool. Samples collected from 6 localities in Cameroon in 2021 and 2022 were also used, with four localities (Nyabessang, Santchou, Mangoum and Nkolondom) situated in the southern, and two localties (Ngaoundere and Gounougou) in the northern part of the country. An assessment of the temporal evolution of the allele was also performed by comparing samples from same location collected at different timepoints notably in Cameroon [Mangoum (2018 vs 2022) and Santchou (2010 vs 2022)].

### Combined impact of CYP6P3 and VGSC knockdown resistance mutation Kdr on insecticide resistance

To assess the combined impact of knockdown resistance (kdr) and CYP6P3 mutations on pyrethroid resistance and on the efficacy of insecticide-treated nets, the L1014F-kdr and CYP6P3-E205D mutations were both genotyped in pyrethroid-exposed (permethrin, deltamethrin and α-cypermethrin) F4 hybrid mosquitoes. In addition, F3 hybrid mosquitoes exposed to insecticide-treated nets were also genotyped. The L1014F-kdr marker was genotyped using the LNA probe-based Taqman assay previously design by Lynd and colleagues (27).

